# A Simple Procedure for Bacterial Expression and Purification of the Fragile X Protein Family

**DOI:** 10.1101/2020.06.24.169243

**Authors:** Madison Edwards, Mingzhi Xu, Simpson Joseph

## Abstract

The fragile X protein family consists of three RNA-binding proteins involved in translational regulation. Fragile X mental retardation protein (FMRP) is well-studied, as its loss leads to fragile X syndrome, a neurodevelopmental disorder which is the most prevalent form of inherited mental retardation and the primary monogenetic cause of autism. Fragile X related proteins 1 and 2 (FXR1P & FXR2P) are autosomal paralogs of FMRP that are involved in promoting muscle development and neural development, respectively. There is great interest in studying this family of proteins, yet researchers have faced much difficulty in expressing and purifying the full-length versions of these proteins in sufficient quantities. We have developed a simple, rapid, and inexpensive procedure that allows for the recombinant expression and purification of full-length human FMRP, FXR1P, and FXR2P from *Escherichia coli* in high yields, free of protein and nucleic acid contamination. In order to assess the proteins’ function after purification, we confirmed their binding to pseudoknot and G-quadruplex forming RNAs.

## Introduction

The fragile X protein (FXP) family consists of three RNA-binding, ribosome-associating proteins involved in translational regulation: fragile X-related protein 1 (FXR1P), fragile X-related protein 2 (FXR2P), and the most well-known, fragile X mental retardation protein (FMRP)^1,2,3,4^. FMRP’s role in translation repression has been studied extensively, as loss of FMRP expression results in a neurodevelopmental disorder called fragile X syndrome (FXS), the most prevalent form of inherited intellectual disability, and the primary monogenic cause of autism spectrum disorders^5,6,7^. FXS predominantly results from a CGG trinucleotide repeat expansion in the 5’ untranslated region of the *FMR1* gene^6,7^. The expanded repeats are hypermethylated causing transcriptional silencing of the *FMR1* gene, leading to a deficiency or absence of FMRP^6,7,8,9^. Patients with this disorder may experience seizures, hyperactivity, anxiety, and poor language development^7^. On a cellular level, patients with FXS possess a greater density of dendritic spines, and increased numbers of long and immature-shaped spines^10^. It is estimated that 1/5,000 males and 1/4,000-8,000 females possess the full FXS mutation^7^.

While perhaps lesser known, FMRP’s autosomal paralogs FXR2P and FXR1P are also of interest for their role in translational regulation^1,2,11^. FXR2P-deficient mice have impaired dendritic maturation of new neurons, with new neurons possessing shorter and less complex dendrites compared to wild-type mice^12^. These mice revealed decreased neural connectivity as new neurons with shorter dendrites connected to fewer presynaptic neurons^12^. Mice deficient in FXR2P displayed atypical gene expression in the brain and altered behavior, such as hyperactivity, reduced sensitivity to heat stimuli, and reduced prepulse inhibition^13,14^.

FXR1P is unique among the fragile X proteins in that three (e-g) of the seven isoforms in mice (a-g) show strong expression in cardiac and/or skeletal muscle^4,15,16,17,18^. In humans FXR1P mRNA likewise demonstrates alternative splicing and is abundant in heart and skeletal muscle tissue^1,15,17,19^. Elimination of *Fxr1* leads to neonatal lethality in mice, while reduced levels of FXR1P lead to shortened life spans and reduced limb musculature^20^. Furthermore, FXR1P expression is altered in myoblasts from patients with facioscapulohumeral muscular dystrophy^17^.

The genes encoding the FXP family are highly homologous through the first 13 exons of FMRP, although FXR1P and FXR2P lack sequences corresponding to exons 11 and 12 of FMRP^21^. After exon 13 of FMRP, the sequences of the three proteins diverge significantly^21^. This suggests that the three proteins likely arose from multiple gene duplications of a common ancestral gene^21^. The amino acid sequences of these proteins display a high similarity over the first 58-70% of their sequences, but a lower similarity thereafter (Figure 1). Additionally, all three proteins possess RNA-binding domains of interest: three well-conserved K homology (KH) domains, and an arginine-glycine-glycine (RGG) motif with poor conservation^11,22,23^. Another noteworthy feature of the fragile X proteins is their C-terminal intrinsically disordered region (IDR) which constitutes ∼30-43% of the entire protein sequence but has lower sequence conservation (Supplementary Figure 1). IDRs are enriched in RNA-binding proteins compared to the entire human proteome and can promote protein aggregation and phase transitions, serve as sites for post-translational modifications or protein-protein interactions, and bind to RNA both specifically and non-specifically^24,25^. The high sequence conservation in the N-termini of the FXP family suggests they exhibit some functional redundancy, while their divergent C-termini likely contribute to their unique functions.

**Figure 1.**
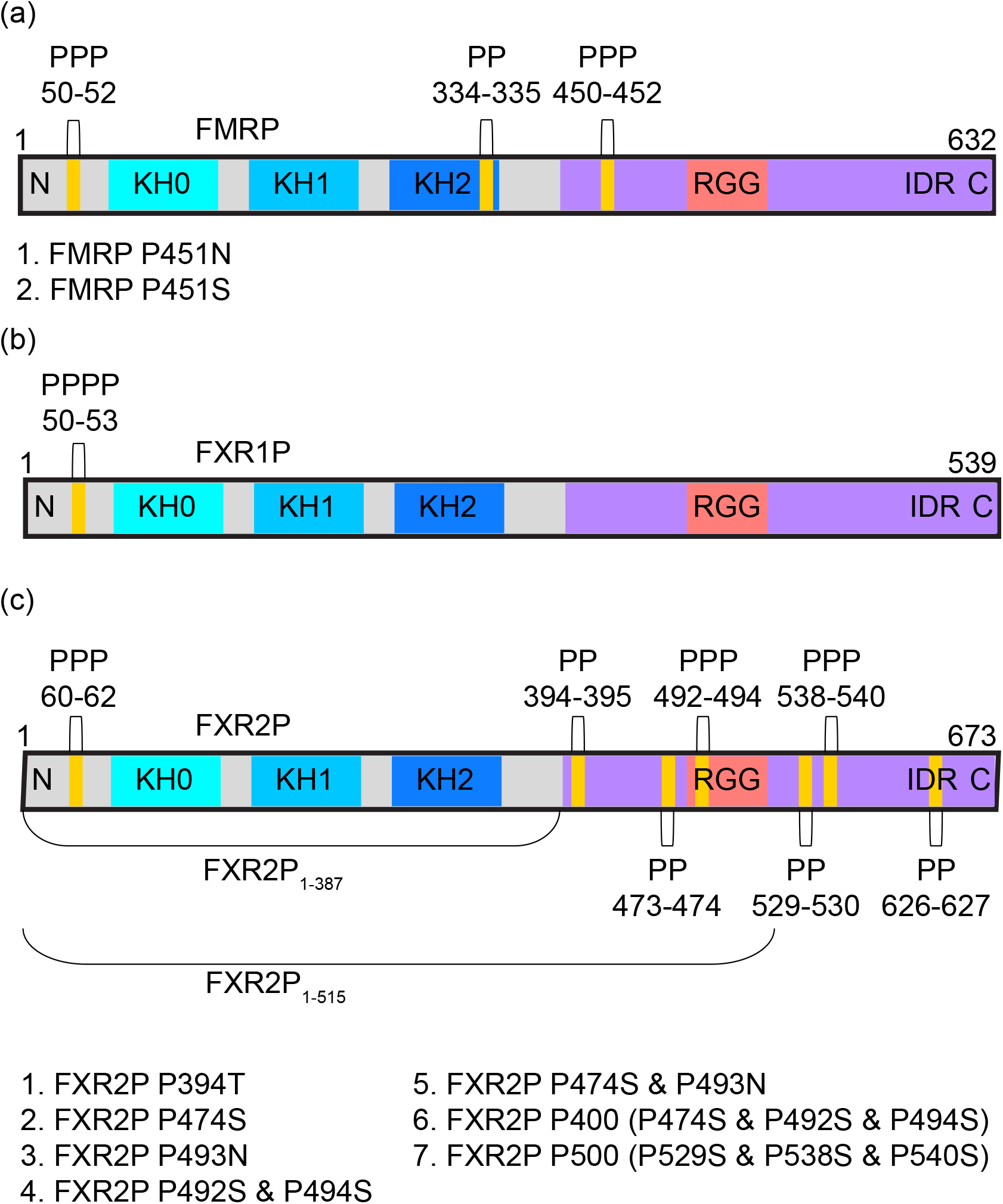
Domains and Diprolyl/Polyproline Stretches of the Fragile X Protein Family. (A) FMRP isoform 1, (B) FXR1P isoform 2, and (C) FXR2P with relevant protein domains and motifs labeled. Diprolyl/polyproline locations are displayed, and shortened FXR2P constructs and FXR2P and FMRP mutants are listed. The Fragile X proteins are well-conserved (72-77% identity) through the sequence RQIG of each protein (located after KH2 domain), but the sequences diverge after this point (31-61% identity). Sequence identities were determined in MUSCLE^59^.

Much emphasis has been placed on proposing mechanisms of translational regulation for the FXP family, with a particular focus on determining their mRNA targets. Many studies have attempted to identify and validate the mRNA targets of FMRP, while several papers have identified targets of FXR1P and FXR2P ^12,26, 27,28,29,30,31,32^. Although there appears to be overlap in the mRNA targets of the FXP family, there is evidence that each protein has unique mRNA targets^11,12, 26,29,31,33^. In order to validate, analyze, or compare the mRNA targets of the FXP family, researchers often test the direct binding of each protein to its mRNA targets in vitro. These studies allow researchers to identify binding sites within a target mRNA or test binding to in vitro selected RNAs, leading to the identification of sequence motifs or structural features the proteins may recognize in vivo^34,35,36^. Such studies have identified G-quadruplexes and kissing complexes as RNA features recognized by FMRP^11,34,35,36^. Thus, it is important to purify these proteins in sufficient quantities, with sufficient purity for in vitro assays.

However, researchers have faced difficulty in purifying full-length fragile X proteins due to their poor expression, the production of truncated proteins (TPs), their tendency to aggregate and precipitate, and their instability in solution when not bound to RNA^31,37,38,39^. In order to overcome such obstacles, researchers have implemented strategies such as plasmids with tRNAs for rare codons to improve expression, extensive and stringent washes to remove contaminant proteins and TPs, purification from inclusion bodies, and purification under denaturing conditions which requires protein refolding and often lengthy dialysis steps^31,38,40^. Many researchers have also purified specific regions or domains instead of purifying the full-length proteins^11,37,40^.

Others have purified the fragile X Proteins from mammalian cells or the SF9/baculovirus system, but these systems are more expensive and time consuming than purifying from *E. coli*, and yields can still be low^27,31,36,40,41,42^. One group noted that when purifying from HEK293 cells, it was challenging to obtain high yields of full-length human FMRP, or to obtain the protein at concentrations above ∼1 μM, noting that this could be due to low expression or a tendency of the protein to precipitate at higher concentrations^43^. Another potential advantage of purification from *E. coli* is that it allows for the purification of the proteins without post-translational modifications, which have been proposed to have a role in the protein’s function^44,45^. Thus, purification from *E. coli* allows researchers to analyze the effects of specific post-translational modifications.

Our purification protocol improves upon previous methods by allowing the FXP family to be purified using a single, simple protocol. This protocol is fast and inexpensive as the proteins are all recombinantly expressed and purified in *E. coli*, and the materials required are available in most biochemistry laboratories. Briefly, mutations were implemented to disrupt ribosomal stalling proline-rich motifs within the protein sequences. These mutations in tandem with a maltose-binding protein (MBP) tag dramatically boosted the expression of the proteins. The mutations also reduced the production of TPs. An ammonium sulfate precipitation step removed the majority of protein contaminants, while the use of a heparin column removed remaining protein contaminants, TPs, and nucleic acid contamination. The final protein samples were pure and obtained in high yields of 1-9 mg from 2 L of culture. Finally, the purified proteins bound to G-quadruplex and kissing complex RNAs, demonstrating that they are functional.

## Results

### Expression of Recombinant Fragile X Proteins

We initially attempted to purify FXR2P from *E. coli* using an N-terminal 6X His-tag, and in doing so, encountered extremely poor expression of the full-length protein. In fact, the primary protein obtained was *E. coli* bifunctional polymyxin resistance protein ArnA, which has a similar molecular weight (74 kDa) to His_6_-FXR2P (75 kDa). Furthermore, ArnA forms a hexamer with surface-exposed patches of histidine residues that bind to nickel beads^46^. To improve recombinant expression, codon optimized sequences were purchased for human FXR2P and FMRP (isoform 1); FXR1P (isoform 2) expression was sufficient, so a codon optimized sequence was not used. All three genes were cloned into standard protein expression vectors and transformed into *E. coli* Rosetta 2(DE3)pLysS for expression tests.

After codon optimization, the expression of FXR2P was still low, so several fusion tags were tested to boost protein expression. Only an N-terminal SUMO or MBP tag seemed to boost the expression of FXR2P, which is supported by the observation that MBP and thioredoxin (TrX) tags are the best N-terminal tags for promoting protein solubility^47^. There is a decreasing likelihood of soluble expression of mammalian proteins as their molecular weight increases, while the presence of low complexity regions within a protein correlates with reduced soluble expression in *E. coli*, which could both explain FXR2P’s poor expression without the MBP tag^47^. Although the His_6_-MBP tag is large (∼44 kDa), in some cases it promotes the proper folding of the attached protein into the biologically active conformation, and in our study, did not contribute to the RNA-binding capabilities of the FXP family^48^. Moreover, it may be possible to cleave the MBP-tag after purification (data not shown). Thus, we selected the MBP tag for our studies.

The MBP tag boosted expression of FXR2P, but we observed the production of many TPs, which we hypothesized were a result of diprolyl and polyproline stretches within FXR2P (Figures 1C and 2C). As the ring structure of proline makes it a poor peptide bond donor and acceptor, two or more consecutive prolines can cause ribosomes to stall during translation^49,50^. Due to nascent chain-mediated stalling of ribosomes, ribosomal rescue mechanisms may release the ribosomes and unfinished proteins from the mRNA chain, leading to the TPs we observe^51,52^.

**Figure 2.**
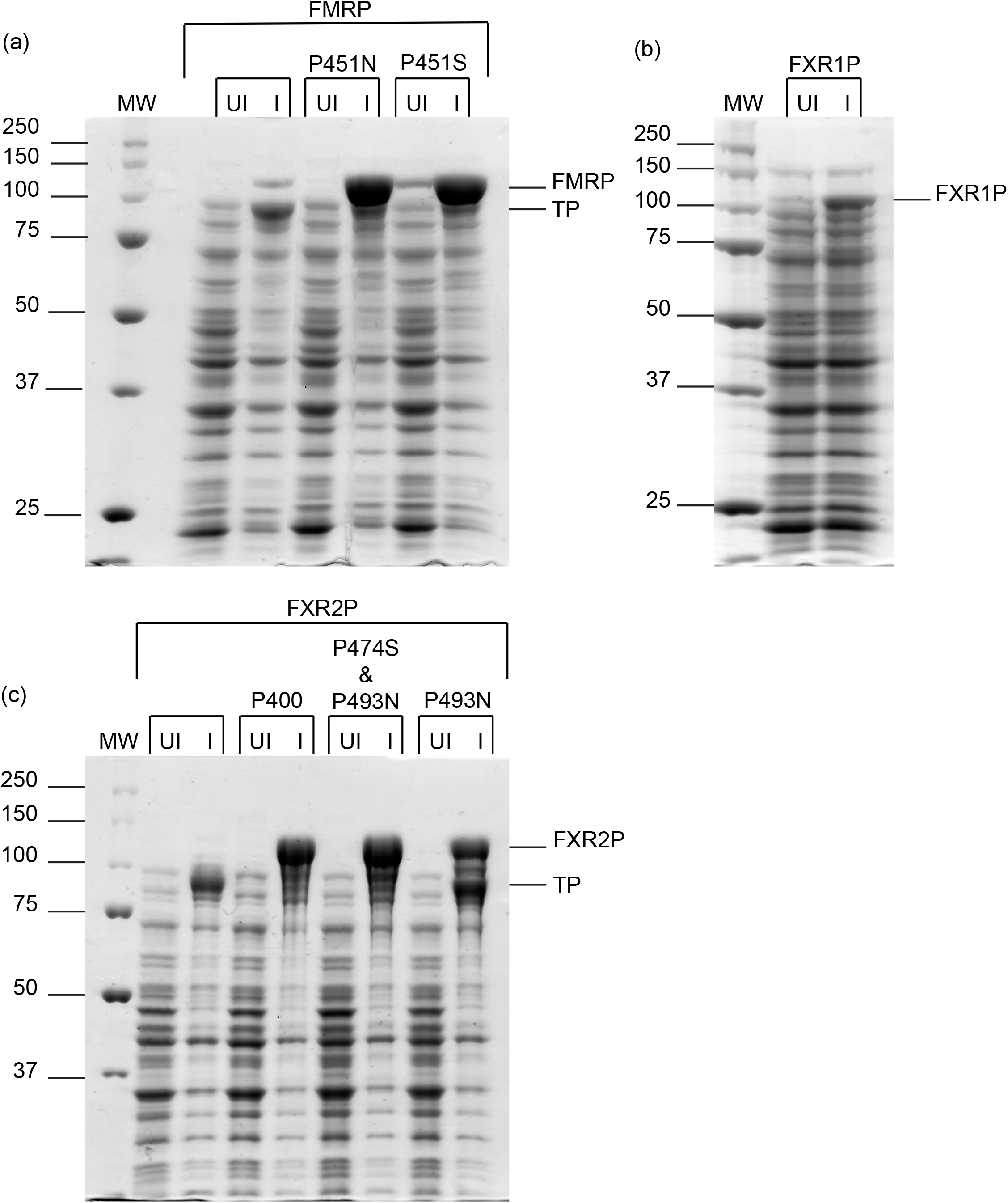
Mutations Enhance the Expression of FMRP and FXR2P. (A) His_6_-MBP-FMRP (∼115 kDa) and mutants, (B) His_6_-MBP-FXR1P (∼104 kDa), and (C) His_6_-MBP-FXR2P (∼117 kDa) and mutants. Uninduced (UI) and induced samples (I) were compared for each construct. FMRP P451S and FMRP P451N mutants exhibited similar increases in expression and reduction in truncated proteins (TP). FXR1P was not mutated, and a non-codon optimized sequence was used, which may explain the lower expression compared to FMRP or FXR2P mutants. FXR2P P474S & P493N was selected as it had higher expression and less TPs than FXR2P P493N, but less mutations than FXR2P P400.

To enhance expression of the full-length protein and reduce the production of TPs we tried two approaches (1) co-expression with elongation factor P (EF-P) which alleviates ribosomal stalling in short proline-rich motifs by stimulating peptide bond formation and (2) mutations to disrupt the diprolyl and polyproline motifs^53^. Although co-expression with EF-P enhanced FXR2P expression, we did not see similar results with FMRP (Supplementary Figure 2). Furthermore, expression with EF-P did not appear to alleviate the production of TPs. In contrast, mutations dramatically boosted expression of full-length FXR2P and FMRP while reducing the production of TPs (Figure 2A & C).

In creating mutations, we aimed to preserve the original protein sequence as much as possible (Supplementary Figure 3). We therefore created and compared the expression of two shortened constructs of FXR2P and seven mutants, which enabled us to determine that the polyproline stretch from residues 492-494 has the greatest contribution towards ribosomal stalling and the production of TPs (Supplementary Figure 4). However, we found we could further enhance expression and reduce TPs by also disrupting the nearby diprolyl motif from residues 473-474. Interestingly, the diprolyl and polyproline stretches at residues 529-530 and 538-540, which are situated closer to one another, do not seem to cause much ribosomal stalling or TPs (see FXR2P P500 mutant). This ultimately led us to select a FXR2P double mutant (FXR2P P474S & P493N) for future purification attempts (Figure 2C).

Based on the results for FXR2P, we decided to mutate the polyproline motif in FMRP that was also located within its disordered region, residues 450-452. Mass spectrometry results of the major TP of FMRP indicated that truncation was occurring after P451, providing further evidence that the TPs are a result of ribosomal stalling at proline-rich motifs (data not shown). Of the two mutants, we selected FMRP P451S for purification attempts (Figure 2A). FXR1P did not need to be mutated as it contains only one polyproline motif, however we did observe a truncation for FXR1P (Figures 1B and 2B). This TP does not appear to be due to ribosomal stalling as mass spectrometry results suggested the truncation occurs within the KH1 domain (data not shown). This TP is lacking the KH2 domain and C-terminal disordered region and appeared to be more stable than full-length FXR1P.

### Purification of the Fragile X Proteins

After selecting a FXR2P and FMRP mutant for purification, we set out to identify a single expression and purification scheme for all three proteins. We grew our cells at 37°C until the OD600 was ∼0.4, then allowed the cells to grow an additional 14-16 hours at ∼14°C after induction with Isopropyl β- d-1-thiogalactopyranoside (IPTG). Initially we attempted amylose resin for protein affinity chromatography, however, we encountered poor binding to the amylose resin unless we implemented a purification step prior to batch binding, and we were unable to remove all the TPs. We therefore switched to an ammonium sulfate precipitation followed by a heparin column. After harvesting and lysing the cells and clarifying the lysate, we were pleased to discover that all three proteins could be precipitated at relatively low percentages of ammonium sulfate, while the majority of *E. coli* proteins remained in the supernatant (Supplementary Figures 5-7). After allowing the fragile X protein to precipitate, the sample was centrifuged, the supernatant removed, and the pellet resuspended. The resulting solution was dialyzed overnight to remove ammonium sulfate as salt must be removed prior to the heparin column. After dialysis, soluble protein was further purified through fast protein liquid chromatography (FPLC) with a heparin column. Using an increasing salt gradient from 0-1 M NaCl (or KCl) we were able to remove nucleic acid contamination, residual protein contaminants, and TPs from the fragile X proteins (Supplementary Figure 8). Interestingly, we observed that the C-terminally truncated His_6_-MBP-FXR1P that is predicted to be missing the KH2 domain and disordered region (∼75 kDa) did not appear to bind to the heparin column and was present predominantly in the flow-through (Supplementary Figure 6). Additionally, the FXR2P TPs began eluting prior to the full-length protein, whereas the full-length was predominantly eluted at high salt concentrations (Supplementary Figure 7). Thus, it appears the fragile X proteins interact with the heparin column through their C-terminal disordered regions. Through the use of a single ammonium sulfate precipitation step and a heparin column, we were able to purify all three fragile X proteins (Figure 3).

**Figure 3.**
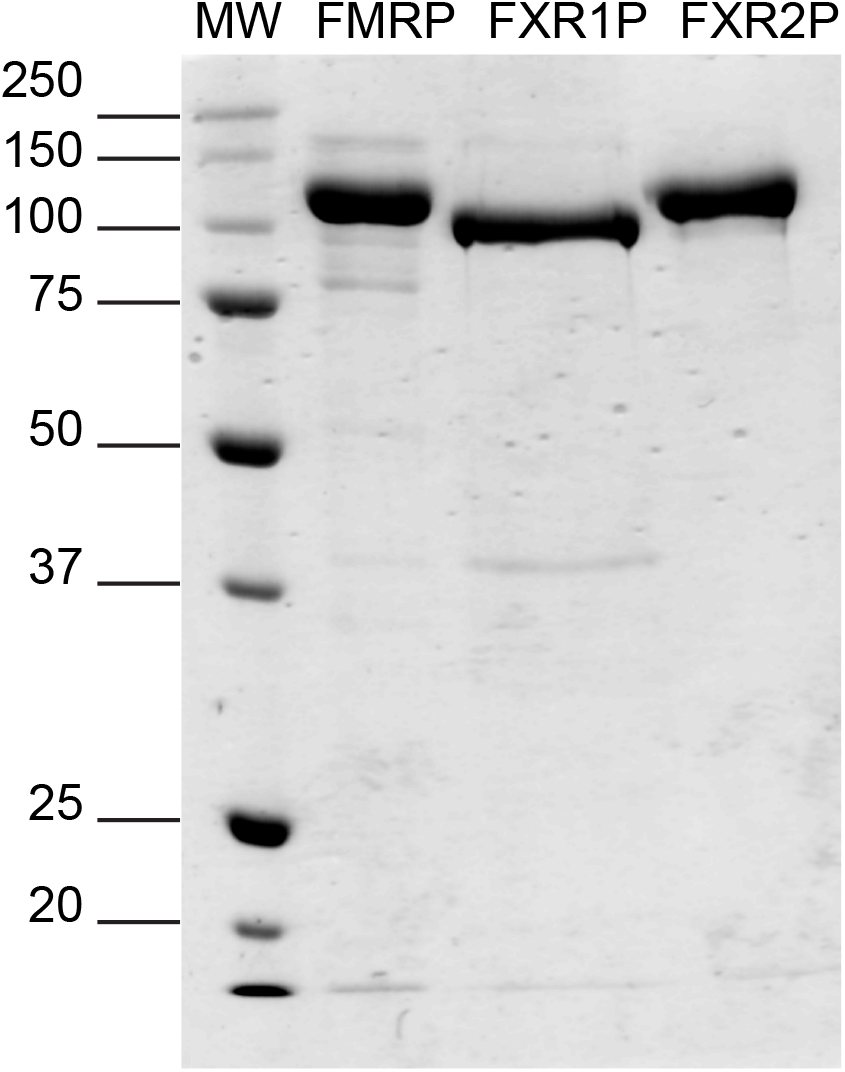
Purified Fragile X Proteins. Approximately 2 μg of each fusion protein was analyzed by sodium dodecyl sulfate polyacrylamide gel electrophoresis (SDS-PAGE). His_6_-MBP-FMRP P451S (∼115 kDa), His_6_-MBP-FXR1P (∼104 kDa), and His_6_-MBP-FXR2P P474S & P493N (∼117 kDa).

### Concentration and Storage of the Fragile X Proteins

After purification we concentrated FMRP and FXR1P by centrifugation. However, we observed a loss in yield, perhaps due to the protein sticking to the concentrator and due to protein precipitation. In our hands, even with a His_6_-MBP tag which promotes solubility, the proteins will precipitate if concentrated too much after removal of nucleic acid contamination^47,48^. FMRP has been concentrated to ∼16 μM but is not stable for long at this concentration and drops to ∼3-6 μM over time. FXR1P and FXR2P precipitate more readily. We were able to concentrate FXR1P to 8 μM but precipitation was visible at ∼3 μM. FXR1P and FXR2P fractions from the heparin column at ∼5-7 μM seem stable whereas fractions at ∼9-13 μM had visible precipitation as they eluted. Once purified from nucleic acids, the fragile X proteins seem stable at ∼6 μM initially and appear to precipitate slowly over time.

To maximize yield, we suggest researchers avoid concentrating, concentrate minimally, or concentrate right before use. When we centrifuged our concentrated protein samples after storage at 4°C or −15°C to remove precipitated protein prior to use, we noticed a decrease in concentration of the samples over time. This is likely due to precipitation. Storage at −80°C may prevent this, although thawing sometimes induces precipitation. The tendencies of the fragile X proteins to form aggregates, precipitate during concentration, or not concentrate past a certain concentration have been noted by other researchers^37,38,39,43^. Concentrating appears to induce aggregation and precipitation, which may have a function in vivo, namely in the proteins’ presence in ribonucleoprotein granules^39^. In rat brain, FXR1P predominantly forms oligomers or insoluble aggregates, while monomers are nearly undetectable^40^. We therefore recommend the MBP tagged fragile X proteins be stored at concentrations of 6 μM or less at 4°C for use within 1-2 weeks or −80°C for long-term storage. We also suggest researchers centrifuge stored samples to remove insoluble aggregates then remeasure protein concentration prior to use.

### Analysis of the RNA-binding Activity of the Fragile X Proteins

The functionality of the purified proteins was verified by testing the FXP family’s binding to a G-quadruplex forming RNA, a well-known target of FMRP, which is bound by the RGG motif, and a target of FXR1P^11,30,35,36,41,54^. We chose to test the proteins’ binding to poly-G17U as our lab has identified poly-G17U as a G-quadruplex forming RNA^55^. For a negative control we used CR1, an RNA with no predicted G-quadruplex forming capability.

As predicted, all proteins showed high affinity binding to poly-G17U, and no binding to CR1 in the concentration range tested (Figure 4A-C). Our fluorescence anisotropy results reinforce the hypothesis that the fragile X proteins have different affinities for mRNA targets: FXR2P showed the greatest affinity for poly-G17U, and FXR1P the least (FXR2P K_d_ = 3.1 ± 0.4, FMRP K_d_ = 5.6 ± 0.6, and FXR1P K_d_ = 11.7 ± 1). To ensure that the His_6_-MBP tag did not contribute to the observed RNA-binding, we tested its binding to poly-G17U and CR1 and observed no binding (Figure 4D).

**Figure 4.**
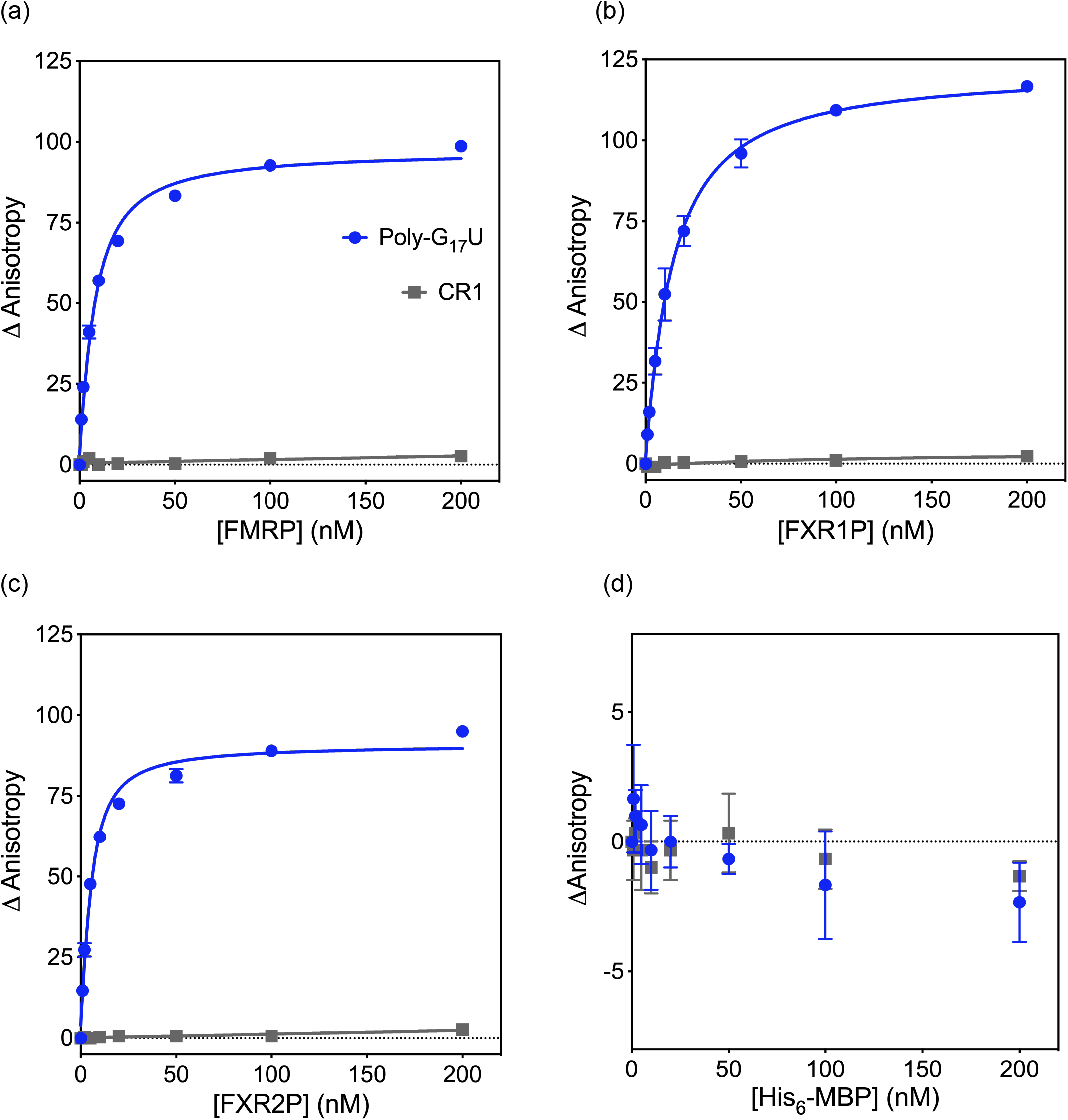
The Fragile X Proteins Bind Poly-G_17_U. The RNA-binding capabilities of (A) FMRP, (B) FXR1P, and (C) FXR2P were assessed by fluorescence anisotropy. All three proteins bound to poly-G_17_U with high affinity: FMRP K_d_ = 5.6 ± 0.6, FXR1P K_d_ = 11.7 ± 1, and FXR2P K_d_ = 3.1 ± 0.4. No binding was observed for CR1 in the concentration range tested. Data are from three individual trials with error bars for the standard deviation displayed (with the exception of FMRP 1 and 2 nM points for which there was only one trial). (D) His_6_-MBP was tested for binding to poly-G_17_U and CR1 through fluorescence anisotropy and shows no RNA-binding capabilities.

After assessing the binding of the fragile X proteins to a G-quadruplex forming RNA, we tested for binding to a loop-loop pseudoknot, or “kissing complex” RNA, ΔKC2, a shortened version of an in vitro selected target of FMRP called KC2^34^. The KH2 domain of FMRP was found to be necessary and sufficient for FMRP binding to KC2^34^. Furthermore, the KH2 domains of FMRP, FXR1P, and FXR2P bind KC2 RNA with equal affinity^11^. As binding of FMRP to KC2 is dependent on the integrity of the KH2 domain, we felt it valuable to assess the FXP family’s ability to bind ΔKC2^11,34^.

Due to the size of our ΔKC2 (72 nucleotides), we tested for binding by electrophoretic mobility shift analysis (EMSA) using CR1 and poly-G_17_U as negative and positive controls, respectively. As predicted, all proteins showed binding to poly-G_17_U and ΔKC2, indicated by the reduction in free RNA upon the addition of protein, but not to CR1 (Figure 5). A discrete shift is not observed for the RNA-protein complex as it appears this complex is stuck in the wells, possibly because the RNA-protein complexes are large. The fragile X proteins appear to bind more tightly to poly-G_17_U than ΔKC2 which is supported by the K_d_ values we determined for poly-G_17_U (∼3-12 nM) and those reported for the binding of the KH2 domains of the fragile X proteins to KC2 (∼40-60 nM)^11^.

**Figure 5.**
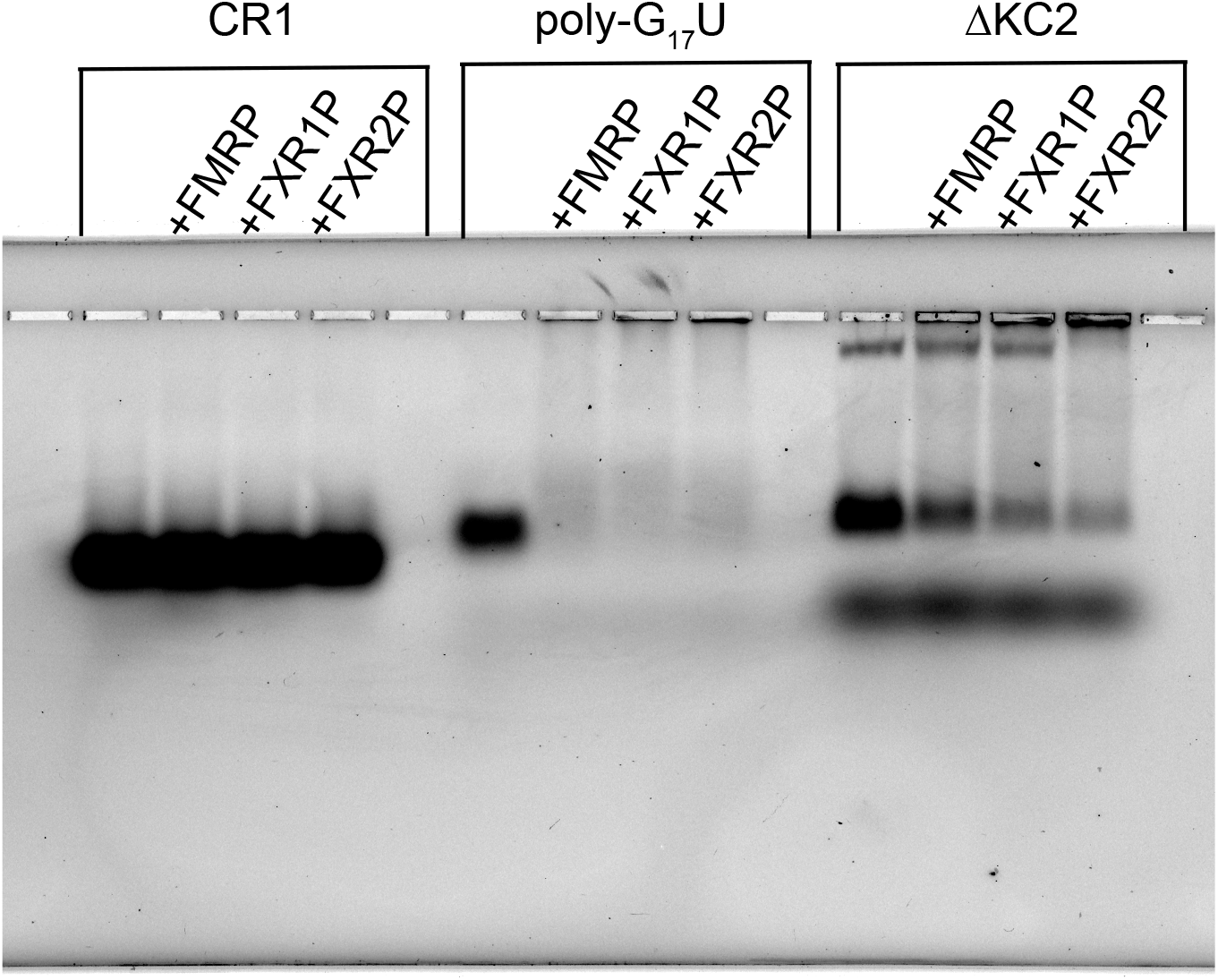
The Fragile X Proteins Bind ΔKC2. Binding was assessed using an agarose EMSA with CR1 and poly-G_17_U included as negative and positive controls, respectively. The bottom band in ΔKC2 lanes is free fluorescein dye.

It is worth mentioning that there are two free RNA species visible in the control lane for ΔKC2. It is likely that the faster migrating species is more compact and folded in the correct confirmation. This is further supported by the observation that the faster migrating species appears to be preferentially bound by FMRP and FXR1P. Darnell et al. also observed two free RNA species for KC2 RNA, and only the faster migrating species was observed to shift with added protein^34^.

## Discussion

Our RNA-binding studies suggest the fragile X proteins are functional. Thus, mutations implemented to boost expression of FMRP and FXR2P do not appear to impact their RNA-binding specificity. We were intrigued by the results of our binding studies to poly-G_17_U, as work by Darnell et. al demonstrated that the C-termini of the fragile X proteins had differing affinity for an in vitro selected G-quadruplex forming RNA (sc1): FMRP bound with high affinity, FXR2P showed lower affinity and non-specific binding, while FXR1P showed no binding^11^. The results we observe may be due to the fact that sc1 was selected using FMRP, while poly-G_17_U may form a generic G-quadruplex structure recognized by all three proteins^35^. Additionally, our results may have been impacted by assessing the binding of the full-length proteins. Future assays testing the RNA-binding specificity of the full-length proteins should yield insightful results as it has been proposed that the multiple RNA-binding domains of the fragile X proteins function cooperatively^26,28^.

Our purification protocol opens the door for compelling research on FXR1P and FXR2P, which have not been studied as extensively as FMRP. As an example, we observed high affinity binding of FXR1P isoform 2 to poly-G_17_U, despite previous results suggesting only the muscle-specific isoforms exhibit high affinity binding for G-quadruplex forming RNAs^41^. Additionally, the ability of FXR2P to bind to G-quadruplexes is not well-documented, yet our results suggest this could be a worthwhile avenue for further research. Our results highlight the utility of our protocol for purifying and comparing the functions of the fragile X proteins in vitro.

In summary, we have identified a rapid, simple, and inexpensive purification protocol for the human FXP family, while many of our techniques can be broadly applied. We found the MBP tag very efficient at enhancing the expression and solubility of our proteins, which may be useful to researchers working with large eukaryotic proteins, or proteins with disordered regions^47,48^. By disrupting ribosomal stalling proline-rich motifs within FMRP and FXR2P we drastically boosted recombinant expression while reducing the production of TPs. This technique, or co-expression with EF-P, may assist in the recombinant expression of eukaryotic proteins, 10% of which possess polyproline motifs^56^. An ammonium sulfate precipitation followed by a heparin column allowed the fragile X proteins to be obtained in high yields, free of *E. coli* protein and nucleic acid contamination. Additionally, this procedure removed the vast majority, and in some cases all, C-terminally TPs. We found the heparin column to be a quick and effective method for removing nucleic acid contamination from nucleic acid binding proteins. All three proteins demonstrated RNA-binding activity through their binding to G-quadruplex and kissing complex forming RNAs. We hope this procedure will mitigate obstacles faced in studying the important roles of the FXP family in translational regulation, and in doing so, promote diverse research questions. Moreover, the techniques described will aid researchers in recombinantly expressing and purifying proteins with poor expression, proline-rich regions, disordered regions, or nucleic acid binding properties.

## Methods

### Creation of Fragile X Protein Expression Vectors

An *E. coli* codon optimized sequence was purchased in pUC57 from GeneWiz for FXR2P, and FMRP was purchased as a gene block from IDT. The human sequence (not optimized for *E. coli*) for FXR1P was purchased from Addgene. The genes coding for the human fragile X proteins (FMRP isoform 1 NCBI Reference Sequence: NP_002015.1, FXR1P isoform 2: NP_001013456.1, and FXR2P NP_004851.2) were introduced through Ligation Independent Cloning into the pMCSG9 vector (DNASU plasmid repository) which provides an N-terminal His_6_-MBP sequence, T7 promoter, ColE1 origin of replication, and ampicillin resistance^57^. For FXR2P only, the TEV protease cleavage site located after the MBP tag was replaced with an HRV 3C protease cleavage site. The resulting plasmids were transformed into DH5α *E. coli* cells (ThermoFisher), purified, and the sequences verified by Sanger sequencing (GeneWiz). After confirming the cloning process, the plasmids were transformed into chemically competent Rosetta 2(DE3)pLysS *E. coli* cells for protein expression (Novagen, chloramphenicol resistance).

#### Primers to insert codon optimized FMRP into pMCSG9

Forward: 5’- TACTTCCAATCCAATGCCATGGAAGAACTGGTGGTTGAAGTGCGTG-3’

Reverse: 5’- TTATCCACTTCCAATGTTACGGCACACCATTGACCAGCGG-3’

#### Primers to insert FXR1P into pMCSG9

Forward: 5’-TACTTCCAATCCAATGCCGCGGAGCTGACGGTGGAGGTT-3’

Reverse: 5’-TTATCCACTTCCAATGTTAATCACATCTTTTGCCTAGCCC-3’

#### Primers to insert codon optimized FXR2P into pMCSG9

Forward: 5’-TACTTCCAATCCAATGCCATGGGCGGTCTGGCGAGC-3’

Reverse: 5’-TTATCCACTTCCAATGTTAGCTCACACCATTCACCATGCTACC-3’

#### Primers to replace TEV site of FXR2P pMCSG9 with an HRV 3C cleavage site

Forward: 5’- CTGGAAGTTCTGTTCCAGGGTCCGATGGGCGGTCTGGCGAGC-3’

Reverse: 5’- GCTACCACCACCACCAGTCTGCGCGTCTTTCAGGG-3’

### Creation of FMRP/FXR2P Mutants and Shortened FXR2P Constructs

Mutations were selected by comparing the amino acid sequence of the human fragile X protein to the same protein in other species in order to avoid mutating highly conserved residues. If possible, prolines were mutated into an amino acid present in another species at the corresponding position (Supplementary Figure 3). Site-directed mutagenesis or sequence deletions were achieved through PCR with designed primers (listed below) on FMRP/FXR2P in pMCSG9. DpnI digestion, PCR purification, T4 PNK treatment, and ligation were performed sequentially after PCR to produce the desired plasmids. The resulting plasmids were transformed into DH5α *E. coli* cells (ThermoFisher), purified, and the sequences verified by Sanger sequencing (GeneWiz). Plasmids containing the desired mutations were transformed into chemically competent Rosetta 2(DE3)pLysS *E. coli* cells (Novagen) to produce desirable cell stocks.

#### Primers to make FXR2P_1-387_

Forward: 5’-TAACATTGGAAGTGGATAACGGATCCG-3’

Reverse: 5’-TTGACGCAGTTGCTCGTCAATC-3’

#### Primers to make FXR2P_1-515_

Forward: 5’-TAACATTGGAAGTGGATAACGGATCCG-3’

Reverse: 5’-ATCCGGGTCTTTCAGCACG-3’

#### Primers to mutate prolines

Codons that introduce a mutation are underlined.

##### FMRP P451S

Forward: 5’ <u>TCT</u>CCGAACCGTACCGATAAAGAAAAGTC 3’

Reverse: 5’ CGGACGAGAGCTTGCACCGATTTG 3’

##### FMRP P451N

Forward: 5’ <u>AAT</u>CCGAACCGTACCGATAAAGAAAAGTC 3’

Reverse: 5’ CGGACGAGAGCTTGCACCGATTTG 3’

##### FXR2P P400 (P474S & P492S & P494S)

Forward: 5’ GACCGGTGGTCGTGGCCGTGGT<u>AGC</u>CCG<u>AGC</u>GCGCCGCGTCCG 3’

Reverse: 5’ GGACGACGACGGCTTTCTTCACCACGGGT<u>GCT</u>CGGATCACGGTCACCC 3’

##### FXR2P P500 (P529S & P538S & P540S)

Forward: 5’ GAGCCGGGCGAA<u>AGC</u>CCG<u>AGC</u>GCGAGCGCGCGTCG 3’

Reverse: 5’ GCTATCCACCGG<u>GCT</u>TTCCGGTTCGCTGGTGTCCAGC 3

##### FXR2P P394T

Forward: 5’- ATTGGCCTGGGTTTTCGT<u>ACC</u>CCGGGTAGCGGCCGTG-3’

Reverse: 5’- TTGACGCAGTTGCTCGTCAATCTGCAGACGCTCCAG-3’

##### FXR2P P474S

Forward: 5’- ACCCGTGGTGAAGAAAGCCGTCG-3’

Reverse: 5’- <u>GCT</u>CGGATCACGGTCACCCGGACC-3’

##### FXR2P P492S & P494S

Forward:5’-CCG<u>AGC</u>GCGCCGCGTCCGACCAGC-3’

Reverse: 5’- <u>GCT</u>ACCACGGCCACGACCACCG-3’

##### FXR2P P493N

Forward: 5’-<u>AAC</u>CCGGCGCCGCGTCCGACCAGCC-3’

Reverse: 5’-CGGACCACGGCCACGACCACCGG-3’

##### FXR2P P474S & P493N

Made by taking FXR2P P493N in pMCSG9 and using the primers for FXR2P P474S to add the second mutation.

### Creation of EF-P and FMRP/FXR2P Co-expression Vectors

A codon optimized sequence for Elongation Factor P (NCBI Reference Sequence: P0A6N4.2) was purchased as His_6_-EF-P in pUC57 (Gene Universal). Seamless cloning was used to insert His_6_-EF-P into a pDSG310 vector (a gift from Ingmar Riedel-Kruse: Addgene plasmid #115611; http://n2t.net/addgene:115611; RRID: Addgene_115611). The pDSG310 vector was selected as it has an arabinose regulated promoter (pBAD), p15A origin of replication, and kanamycin resistance, which are all distinct from those of the pMCSG9 vector used for FMRP and FXR2P^58^. Using primers that enabled seamless cloning, PCR was used to prepare the His_6_-EF-P DNA for insertion into pDSG310, and the backbone of pDSG310 was likewise prepared. The insert and pDSG310 backbone were digested with BbsI and ligated, and the ligated plasmid was transformed into DH5α *E. coli* cells which produced colonies containing viable EF-P containing plasmids. The resulting plasmids were purified, and the sequences verified by Sanger sequencing (GeneWiz). To make cells co-expressing EF-P and FMRP or FXR2P, chemically competent Rosetta 2(DE3)pLysS *E. coli* cells (Novagen) were transformed with 1:1 EF-P plasmid: FMRP/FXR2P plasmid (200 ng each). A control cell stock containing only EF-P in Rosetta 2(DE3)pLysS *E. coli* cells was also produced.

#### Primers to insert EF-P into pDSG310

Forward: 5’ CGTCGAGAAGACTACTAGATGCACCATCATCATCATC 3’

Reverse: 5’ CGTCGAGAAGACTTATTTCACGCGGCTCACATATTC 3’

#### Primers to linearize pDSG310

Forward: 5’ CGTCGAGAAGACTGAAATAATAATACTAGAGCCAGGCATCAAATAAAAC 3’

Reverse: 5’ CGTCGAGAAGACATCTAGTATTTCTCCTCTTTCTCTAGTAGCTAGC 3’

### Expression Tests of Fragile X Proteins, Mutants, & Co-expression with EF-P

Overnight cultures containing 100 μg/mL ampicillin and 25 μg/mL chloramphenicol (these are the concentrations of antibiotics used in all cultures) were inoculated with the appropriate Rosetta 2(DE3)pLysS *E. coli* cells from a glycerol stock. For co-expression tests with EF-P, kanamycin was also added to a final concentration of 50 μg/mL. Three milliliter overnight cultures were incubated ∼16-20 hours at 37°C, ∼215 RPM. The following day, LB broth containing ampicillin and chloramphenicol (and kanamycin for cells co-expressing EF-P) was inoculated with the corresponding overnight culture at a ratio of 0.005:1 overnight culture: LB broth. The cultures were incubated at 37°C, ∼215 RPM until the OD600 reached ∼0.4-0.6, although an OD600 of up to 0.8 was allowed in some cases. At this time, the cultures were split into equal volumes (3 mL each) to create an uninduced and induced sample. To the induced samples, Isopropyl β- d-1-thiogalactopyranoside (IPTG) was added to a final concentration of 0.4 mM. For co-expression with EF-P, arabinose was also added to a final concentration of 0.1% to induce the expression of EF-P. The samples were then incubated for an additional 3 hours at 37°C, ∼215 RPM (2 hours for FMRP and mutants Figure 2). For the initial testing of FXR2P mutants only, (Supplementary Figure 4) expression at 14 °C for ∼18 hours was performed. These cultures were cooled for 10 min at ∼0°C to slow cell growth prior to the addition of IPTG.

After expression, samples were prepared for analysis by sodium dodecyl sulfate–polyacrylamide gel electrophoresis (SDS-PAGE). The OD600 of a ¼th dilution of cultures were obtained and used to determine the OD600 of the stock solution. For each culture, 500 μL of sample was centrifuged at 16,100 RCF to pellet the cells. After removing the supernatant, the pellet was resuspended in 25 μL of resuspension buffer (10 mM Tris pH 7.5, 20 mM NaCl) per 0.5 OD600 unit. After resuspending, 60 μL of the cell pellet resuspension was combined with 15 μL of 5X SDS-PAGE loading dye. The samples were boiled at ∼95 °C for 10 minutes, then spun down for a few seconds. The samples were mixed, and 10 μL of each sample was loaded onto a 10% SDS polyacrylamide gel that was run for 10 min at 100V, followed by 180V until the dye front ran off the gel (∼55 minutes). The protein bands were visualized by staining the gels with Coomassie Brilliant Blue.

Comparison of full-length FXR2P expression with and without EF-P co-expression was performed by analyzing band intensity for full-length FXR2P in ImageJ. Fold change of expression was calculated by normalizing the band intensity of full-length FXR2P when co-expressed with EF-P to the band intensity of full-length FXR2P without EF-P expression. Co-expression with EF-P led to a 1.93 ± 0.27-fold increase in FXR2P expression; error reported is the standard deviation.

### Expression and Purification of Recombinant Fragile X Proteins

For each purification, two 4 L flasks were prepared with 1 L of LB broth with ampicillin and chloramphenicol (2 liters of cell culture), and each flask was inoculated with 5 mL of an overnight culture (0.005:1 overnight culture: LB broth). The cells were incubated at 37°C, ∼215 RPM until the OD_600_ reached ∼ 0.4-0.5; this step generally took 4-5 hours. The cultures were then cooled for 20 minutes by transferring to an incubator at 14°C, ∼150 RPM. The expression of each protein was induced with the addition of IPTG to a final concentration of 0.4 mM. The induction was carried out for 13-15 hours at 14°C, ∼150 RPM.

After induction the cells were split into six 500 mL centrifuge flasks and pelleted by centrifuging at 4,420 RCF, 4 °C, 15 min (Beckman J2-HC Centrifuge). Pellets were transferred to a 50 mL polypropylene Falcon tube and weighed; the typical weight was 6.4-7.4 grams from 2 L of culture. The pellet was then resuspended in lysis buffer with no salt (50 mM Tris pH 7.5, 1 mM ethylenediaminetetraacetic acid (EDTA), 1 mM dithiothreitol (DTT), and 1 mM phenylmethylsulfonyl fluoride (PMSF)) to a total volume of ∼50 mL. The resuspended cells were sonicated (Branson Digital Sonifier) on ice with ten 8 second pulses at an amplitude of 60% interspersed with 1 min pauses. The crude lysate was then clarified by centrifuging at 50,271 RCF, 30 min, 4 °C (Beckman Coulter Optima LE-80K Ultracentrifuge). The clarified lysate was placed in a beaker on a stir plate at 4 °C and concentrated ammonium sulfate (5 mM HEPES pH 7.5) at 4 °C (concentration is temperature dependent, but ∼3.8 M at 0°C) was added to a final concentration of 20% for FXR1P and FXR2P and 25% for FMRP.

After ammonium sulfate addition, the clarified lysate was allowed to sit at 4°C for at least an hour. Subsequently, the solution was centrifuged at 5,087 RCF, 15 min, 4°C (Sigma 4K15C) and the pellet was resuspended in 60-75 mL of lysis buffer. The resuspension was dialyzed in 2L of dialysis buffer (50 mM Tris pH 7.5, 1 mM EDTA, and 1 mM DTT) for at least 16 hours. After dialyzing, the protein solution was centrifuged at 5,087 RCF, 30 min, 4°C in a swinging bucket centrifuge (Sigma 4K15C) to remove insoluble protein.

Fifty milliliters of the supernatant was loaded onto a 50 mL superloop (Amersham Biosciences) and bound to a 5 mL heparin column (HiTrap Heparin HP 1X5 mL, GE Healthcare) by fast protein liquid chromatography (ÄKTApurifier, Amersham Pharmacia Biotech) that had been pre-equilibrated with at least 5 column volumes (CV) of 0 M salt buffer (25 mM Tris pH 7.5, 1 mM EDTA, 1 mM DTT, 25% glycerol). The typical mass of protein loaded onto the column ranged from 17-130 mg. After collecting the flow-through the column was washed with 2 CV of 0 M salt buffer. The proteins were subsequently eluted with a constant salt gradient that started with 0 M salt buffer (25 mM Tris pH 7.5, 1 mM EDTA, 1 mM DTT, 25% glycerol) and increased the salt concentration over 30 CV, ending with 1 M salt buffer (25 mM Tris pH 7.5, 1 mM EDTA, 1 mM DTT, 25% glycerol, 1 M NaCl). Each of the fragile X proteins eluted within a unique range of salt concentrations. Pure FMRP fractions eluted from ∼500-600 mM NaCl, with the peak max at ∼560 mM, FXR1P at ∼560-700 mM NaCl, peak max at ∼640 mM, and FXR2P over a large range, however the most full-length with the least TPs eluted from ∼700-830 mM NaCl with the peak max at ∼730 mM (Supplementary Figures 5-7). It is important to note that KCl can be used instead of NaCl as we successfully purified FXR1P with KCl, which led the protein to elute at lower salt concentrations ∼460-600 mM, peak max at ∼520 mM.

After analyzing the elution fractions by SDS-PAGE, the desired fractions were either pooled and concentrated by centrifugation at 2,493 RCF, 4°C through a 15 mL 50 kDa MW cutoff concentrator (Amicon Ultra −15 Centrifugal Filters, Millipore Sigma) or individual elution fractions from the heparin column were stored. In either case, the final sample(s) was centrifuged at 16,100 RCF for 10 min at 4°C immediately prior to concentration measurements and storage to remove any precipitated protein. The concentration of the supernatant was determined from the absorbance at 280 nM (A280 values) (Thermo Scientific NanoDrop 2000/2000c spectrophotometer). The pure fragile X proteins were then stored at 4°C, −15°C, or −80°C. The best temperature for storing the proteins appears to be 4°C for short-term storage (1-2 weeks), as partial freezing that can occur at −15°C appeared to induce precipitation. Freezing at −80°C then thawing before use is recommended for long-term storage, although precipitation may occur upon thawing. We therefore recommend centrifuging to remove precipitated protein and remeasuring protein concentration prior to use in assays.

The A260/280 ratio of stored samples is typically ∼0.53-0.62, indicating the nucleic acid contamination has been removed (https://www.biotek.com/resources/docs/PowerWave200_Nucleic_Acid_Purity_Assessment.pdf). Protein yield ranged from ∼1.30-8.59 mg; lower yields around 1-2 mg occurred when the fractions from the heparin column were pooled and concentrated. A loss in concentration occurs during concentration steps as discussed previously. The yield reflects the mass of protein at the end of the purification after removing precipitated protein. Storage buffers for use as blanks for concentration readings and for use in RNA-binding assays were created by mixing 0 M and 1 M salt buffers to create buffers with salt concentrations matching that of the final stored protein samples. The concentration of salt in the protein sample was determined from the elution plots from the FPLC machine.

### G-quadruplex RNA binding of the Fragile X Proteins

To confirm that the fragile X proteins were functional, we tested their binding to G-quadruplex forming RNA through fluorescence anisotropy. To ensure the RNA-binding we observed was not due to the tag we used, we also assessed the binding of His_6_-MBP.

The purified fragile X proteins were centrifuged at 16,100 RCF, 10 min, 4°C with a benchtop centrifuge to remove any precipitated protein prior to each experiment. The supernatants containing soluble protein were obtained and the concentration of protein determined using A280 readings (Thermo Scientific NanoDrop 2000/2000c spectrophotometer). Poly-G_17_U and CR1 labeled with a 3’ fluorescein (Dharmacon) were diluted to 5X concentrations (∼25 nM) and these solutions were kept in the dark during the experiment. The RNAs in these 5X solutions were renatured in renaturation buffer (50 mM Tris pH 7.5, 75 mM KCl, 2 mM MgCl_2_) by heating at 68 °C for 5 minutes, then slow cooled from 68 °C to ∼28°C for ∼1 hour in a water bath. Water, binding buffer, protein storage buffer, protein, and the 5X RNA solution were added in the order listed and mixed together for a final reaction volume of 200 μL. The final reactions contained 20 mM Tris pH 7.5, 75 mM KCl, 5 mM MgCl_2_, 1 μM BSA, 1 mM DTT, 100 ng/μL tRNA (to prevent nonspecific binding), and ∼5 nM RNA. The protein concentrations tested were 0, 1, 2, 5, 10, 20, 50, 100, and 200 nM (for FMRP, only one trial for 1 and 2 nM points). It is important to note that for each protein concentration tested the total volume of protein + protein storage buffer remained constant. In each trial the binding buffer was adjusted to account for the Tris pH 7.5 and DTT that were contributed from the protein storage buffer. For trials with His_6_-MBP we assumed the storage buffer contributions were negligible since the protein was diluted in fragile X protein storage buffer prior to use, and the fragile X protein storage buffer was used as the protein storage buffer in the anisotropy reactions. Reactions were thoroughly mixed and incubated in the dark at room temperature for 1 hour. After incubation, each reaction was added into a 96-well plate for fluorescence anisotropy using a multimode microplate reader (SPARK TECAN). Samples were excited at 485 nm and emission was measured at 535 nm. To determine binding affinities, the anisotropy data from each binding assay were normalized to initial values without protein, plotted, and fit to a quadratic equation as previously described^55^. Three independent trials were performed (except for FMRP 1 and 2 nM points) to determine standard deviations.

#### RNA sequences

18 nucleotides CR1: 5’-GCUAUCCAGAUUCUGAUU-Fluorescein-3’

18 nucleotides poly-G17U: 5’-GGGGGGGGGGGGGGGGGU-Fluorescein-3’

### In Vitro Transcription and Fluorescein Labeling of ΔKC2 RNA

The sequence for ΔKC2 was PCR amplified from a pGEM3Z plasmid (a gift from Eileen Chen) which contains a T7 promoter sequence. Nine 100 μL transcription reactions were set up with 90 μL of the PCR-generated DNA template, 4 mM NTPs, 1X transcription buffer (40 mM Tris pH 8.0, 20 mM MgCl_2_, 2 mM Spermidine, 0.1 % Triton X-100), 5 mM DTT, and ∼0.27 μg of T7 RNA polymerase. Each reaction was treated with 2 units of RQ1 DNase (Promega) for 30 min at 37°C, followed by gel purification on a 10% denaturing polyacrylamide gel. It is important to note that our ΔKC2 RNA contains 10 extra nucleotides at the 5’ end from cloning into pGEM3Z relative to the sequence used by Darnell et al^34^.

To label the RNA, 0.5 nmoles of RNA was 3’ oxidized for 90 minutes at room temperature (0.5 mM KIO4, 100 mM NaOAc pH 5.2) then incubated with fluorescein 5-thiosemicarbizide (FTSC) at 4°C overnight (100 mM NaOAc pH 5.2, 1.5 mM FTSC). The RNA was then purified using a Monarch RNA Clean-up Kit (New England BioLabs).

#### Primers to PCR amplify ΔKC2 RNA

Forward: 5’-GCAACTGTTGGGAAGGGCGATCG-3’

Reverse: 5’-AGACGCACATACCAGCCGCTAGC-3’

### RNA-binding of Fragile X proteins by Electrophoretic Mobility Shift Assay

To confirm the fragile X proteins were functional, we tested their binding to ΔKC2 RNA through an electrophoretic mobility shift assay. Based on the results from fluorescence anisotropy, we used poly-G_17_U and CR1 RNAs as positive and negative controls respectively.

The purified fragile X proteins were centrifuged at 16,100 RCF, 10 min, 4°C with a benchtop centrifuge to remove any precipitated protein prior to each experiment. The supernatants containing soluble protein were obtained and the concentration of protein determined using A280 readings (Thermo Scientific NanoDrop 2000/2000c spectrophotometer). Fluorescein-labeled Poly-G_17_U, CR1, and ΔKC2 were diluted to 10X concentrations (1 μM) and these solutions were kept in the dark during the experiment. The 10X RNA solutions were renatured in renaturation buffer (50 mM Tris pH 7.5, 100 mM KCl, 5 mM MgCl_2_) by heating at 68 °C for 5 minutes, then slow cooled from 68 °C to ∼28°C for ∼1 hour in a water bath. Water, 10X binding buffer, protein storage buffer, protein, and the 10X RNA solution were added in the order listed and mixed together for a final reaction volume of 26 μL. The final reactions contained 50 mM Tris pH 7.5, 145 mM KCl, 5 mM MgCl_2_, 1 μM BSA, 10 mM DTT, 50 ng/μL tRNA (to prevent non-specific binding), ∼100 nM fluorescein-labeled RNA, and for reactions containing protein, 250 nM of protein. It is important to note that for reactions containing FMRP only, which was stored in NaCl, the reactions contained 50 mM KCl + 96 mM NaCl instead of 145 mM KCl. For each protein concentration tested the total volume of protein + protein storage buffer remained constant. In each reaction the binding buffer was adjusted to account for the Tris pH 7.5, KCl, and DTT that were contributed from the protein storage buffer. The reactions were thoroughly mixed and incubated in the dark at room temperature for 1 hour. After incubation, 3 μL of loading dye (xylene cyanol in 50% glycerol) was added to each reaction. A 0.8% agarose gel (SeaKem GTG agarose) was prepared in 0.5X TBM buffer (45 mM Tris pH 8.3, 45 mM borate, 2.5 mM MgCl_2_). After loading 13 μL of each sample, the gel was run at 4°C for 2 hours and 15 minutes at 66 V. The gel was then scanned using a laser scanner (Typhoon FLA 9500, GE Healthcare) and the gel was analyzed in ImageJ.

#### RNA sequences

18 nucleotides CR1: 5’-GCUAUCCAGAUUCUGAUU-Fluorescein-3’

18 nucleotides poly-G17U: 5’-GGGGGGGGGGGGGGGGGU-Fluorescein-3’

72 nucleotides ΔKC2: 5’-GGGCGAAUUCGGGAUUCCGACCAGAAGGGGCUAAGGAAUGGUGGGACGAGCUAGCGGCUGGUAUGUGCGUCU-Fluorescein-3’

## Acknowledgements

We thank Reta Sarsam for cloning FXR1P into pMCSG9 and providing purified His_6_-MBP for RNA-binding assays, Youssi Athar for cloning FMRP into pMCSG9, creating the FMRP mutant plasmids, and sharing the mass spectrometry results of the truncated FMRP species, and Eileen Chen for cloning ΔKC2 into the pGEM3Z plasmid. This work was supported by the National Institutes of Health (R01GM114261 to S.J.) and the Cell and Molecular Genetics Training Program funded by the National Institutes of Health (T32GM007240).

## Author Contributions

M.E. and S.J. designed the experiments with input from M.X. M.E. and M.X. performed the experiments and all authors discussed the results. M.E. and M.X wrote the paper with input from S.J. S.J. supervised all aspects of the work.

