## Supplemental information for "A Simple Procedure for Bacterial Expression and Purification of the Fragile X Protein Family"

**Supplementary Information**

**Supplementary Figure 1**. The Fragile X Proteins Possess Disordered C-termini. Fragile X protein disordered region predictions from IUPred2A using long disorder settings for (A) human FMRP isoform 1, (B) human FXR1P isoform 2, and (C) human FXR2P[59].

**Supplementary Figure 2**. Co-expression with EF-P Enhances Expression of Full-length FXR2P. Co-expression with EF-P (~21 kDa) does not appear to enhance (A) FMRP (~115 kDa) expression but appears to enhance (B) FXR2P (~117 kDa) expression. Arabinose (A) was used to induce EF-P expression, and IPTG (I) to induce FXR2P or FMRP expression. Lanes show the comparison between uninduced (UI), arabinose only (A), IPTG only (I), or samples induced with arabinose and IPTG (A + I). Uninduced and IPTG only conditions are shown for cells containing the EF-P plasmid (FXR2P/FMRP + EF-P), and cells without (FXR2P/FMRP). (C) Comparison of full-length FXR2P expression with and without EF-P co-expression. Expression was determined in ImageJ from the band intensity for full-length FXR2P. Fold change of expression was calculated by normalizing the band intensity of FXR2P when co-expressed with EF-P to the band intensity of control FXR2P. Co-expression with EF-P lead to a $1.93 \pm 0.27$-fold increase in FXR2P expression. Error and error bars represent the standard deviation.

**Supplementary Figure 3**. Multiple sequence alignments of (A) FMRP, (B) FXR1P, and (C) FXR2P from multiple organisms, performed in MUSCLE[26]. Alignments of regions containing polyproline motifs are displayed, with the numbering of polyproline motifs referring to the position within the human sequence.

**Supplementary Figure 4.** Mutating Proline-rich Regions Enhances Full-length FXR2P Expression. Expression test of (A) shortened FXR2P constructs and (B-C) FXR2P mutants used to determine which consecutive prolines were causing ribosomal stalling. Lanes show the comparison between uninduced (UI) and samples induced with IPTG (I). Full-length $His_6$-MBP-FXR2P and mutants are ~117 kDa. $FXR2P_{1-515}$ is ~100 kDa and $FXR2P_{1-387}$ is ~87 kDa. Removing amino acids 388-673 of FXR2P ($FXR2P_{1-387}$) or mutating the polyproline stretch from 492-494 (FXR2P P492S & P494S) increases protein expression.

**Supplementary Figure 5.** FMRP Purification. (A) $His_6$-MBP-FMRP P451S (~115 kDa) is obtained in the lysate and pelleted from *E. coli* proteins at 25% $(NH_4)_2SO_4$. (B) A heparin column removes nucleic acid contamination and the majority of residual *E.coli* and truncated proteins.

**Supplementary Figure 6.** FXR1P Purification. (A) $His_6$-MBP-FXR1P (~104 kDa) is obtained in the lysate and pelleted from *E. coli* proteins at 20% $(NH_4)_2SO_4$. (B-C) A heparin column removes nucleic acid contamination and the majority of residual *E. coli* and truncated proteins. (B) The FXR1P truncated protein (~75 kDa) does not appear to bind to the column and is found predominantly in the flow-through.

**Supplementary Figure 7.** FXR2P Purification. (A) $His_6$-MBP-FXR2P P4742 & P493N (~117 kDa) is obtained in the lysate and pelleted from *E. coli* proteins at 20% $(NH_4)_2SO_4$. (B-C) A heparin column removes nucleic acid contamination and the majority of residual *E. coli* and truncated proteins. Truncated proteins elute before full-length FXR2P. (D) Diluted FXR2P elution fractions reveal that fractions eluted at higher salt concentrations contain less truncated proteins relative to full-length FXR2P.

**Supplementary Figure 8.** High Salt Concentrations Elute the Fragile X Proteins. Heparin column elution fractions for the fragile X proteins displaying A280 absorbance and NaCl concentration. *E. coli* protein contaminants elute at lower salt concentrations than the fragile X proteins. (A) FMRP elutes at ~500-600 mM NaCl, with the peak at ~560 mM NaCl, (B) FXR1P elutes at ~560-700 mM NaCl, with a peak at ~640 mM NaCl, and (C) FXR2P elutes over a large range, however the most full-length with the least truncated proteins eluted from ~700-830 mM NaCl with the peak at ~730 mM.

Supplementary Figure 1

(a)

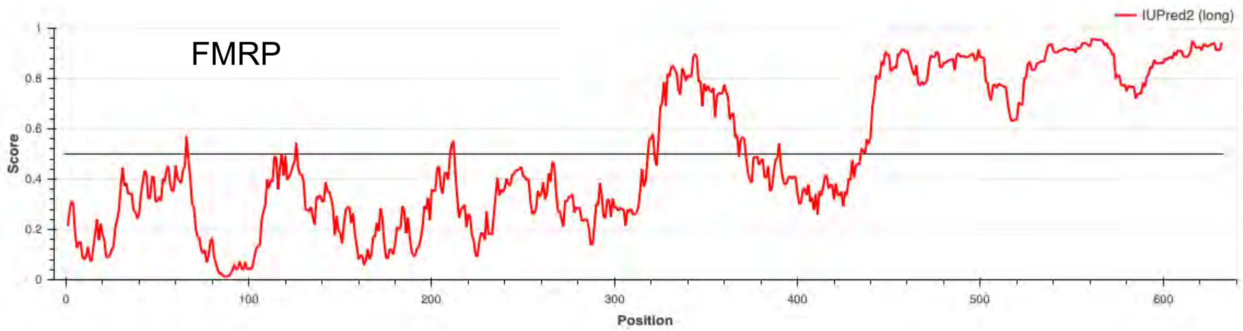


(b)

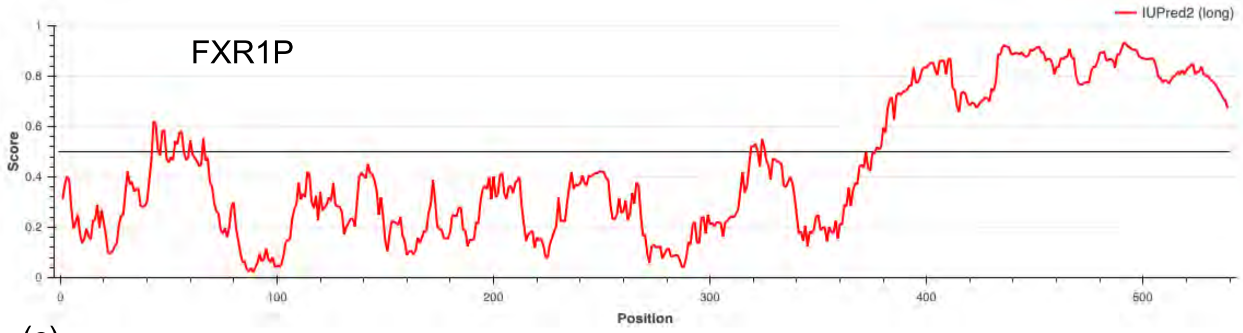


(c)

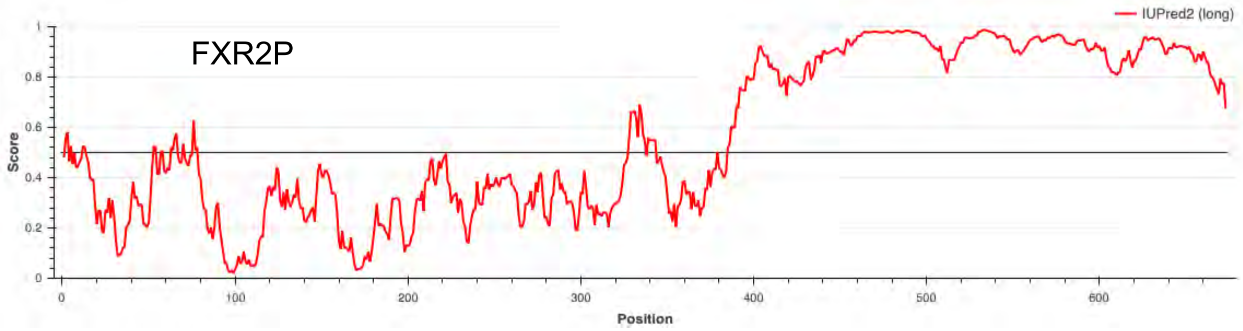

Supplementary Figure 2

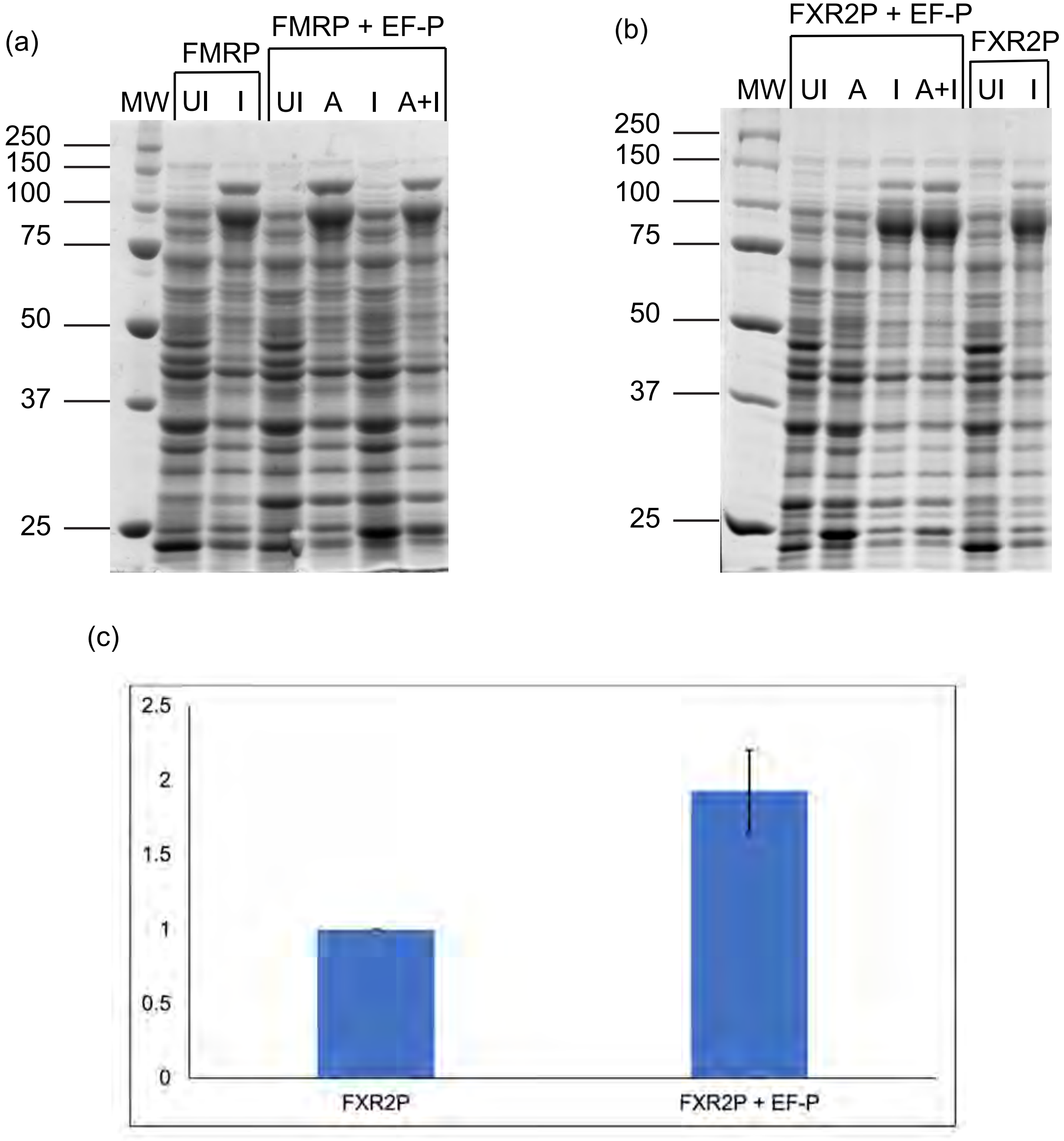

Supplementary Figure 3

(a)

```
                                                                              50-52
FMRPzebrafish      1 MDELAVEVRGASGAFYKAYIKDVHESTVTVAFENNWQPERQISFQDVRFPPPTGFQKEIN 60
FMRPopossum        1 MEELVVEVRGSNGAFYKAFVKDVHEDSITVAFENNWQPERQIPFHDVRFPPPTGYNKDIN 60
FMRPmouse          1 MEELVVEVRGSNGAFYKAFVKDVHEDSITVAFENNWQPERQIPFHDVRFPPPVGYNKDIN 60
FMRPchimpanzee     1 MEELVVEVRGSNGAFYKAFVKDVHEDSITVAFENNWQPDRQIPFHDVRFPPPVGYNKDIN 60
FMRPhuman          1 MEELVVEVRGSNGAFYKAFVKDVHEDSITVAFENNWQPDRQIPFHDVRFPPPVGYNKDIN 60
                     *:**.*****:.******::*****.::**********:***.*:*******.*::*:**
```

```
                                                    334-335
FMRPzebrafish    301GKLIQEVVDKSGVVRVRVEPENDKK---------PSPLE--------------------- 330
FMRPopossum      301GKLIQEIVDKSGVVRVRIEAENDKNIPQEEEIIPQNPLPSNNSRVGSNSSEEKKYLDLKE 360
FMRPmouse        301GKLIQEIVDKSGVVRVRIEAENEKSVPQEEEIMPPSSLPSNNSRVGPNSSEEKKHLDTKE 360
FMRPchimpanzee   301GKLIQEIVDKSGVVRVRIEAENEKNVPQEEEIMPPNSLPSNNSRVGPNAPEEKKHLDIKE 360
FMRPhuman        301GKLIQEIVDKSGVVRVRIEAENEKNVPQEEEIMPPNSLPSNNSRVGPNAPEEKKHLDIKE 360
                    ******:**********:*.**:*.         ..*
```

```
                                                   450-452
FMRPzebrafish    356 LNYLKEVDQLRMERLQIDEQLRQIGGGPRALPGRPEKEKSFMADNGMGPSRGGGKPFGRG 415
FMRPopossum      421 LNYLKEVDQLRLERLQIDEQLRQIGASSRPPPNRADKDKGYMTEDGPGLGR-GSRPYRNR 479
FMRPmouse        420 LNYLKEVDQLRLERLQIDEQLRQIGASSRPPPNRTDKEKGYVTDDGQGMGR-GSRPYRNR 478
FMRPchimpanzee   421 LNYLKEVDQLRLERLQIDEQLRQIGASSRPPPNRTDKEKSYVTDDGQGMGR-GSRPYRNR 479
FMRPhuman        421 LNYLKEVDQLRLERLQIDEQLRQIGASSRPPPNRTDKEKSYVTDDGQGMGR-GSRPYRNR 479
                     ***********:*************...*. *.*.:*:*.:::::* * .* *..*: .
```

(b)

```
                                                                               50-53
FXR1Pzebrafish     1 MEELTVEVRGSNGAYYKGFVRDVHDDSLSISFENNWQPERQVPFSDVRLPPSADTKKEIG 60
FXR1Popossum       0 ------------------------------------------------------------ 0
FXR1Pmouse         1 MAELTVEVRGSNGAFYKGFIKDVHEDSLTVVFENNWQPERQVPFNEVRLPPPPDIKKEIS 60
FXR1Pchimpanzee    1 MAELTVEVRGSNGAFYKGFIKDVHEDSLTVVFENNWQPERQVPFNEVRLPPPPDIKKEIS 60
FXR1Phuman         1 MAELTVEVRGSNGAFYKGFIKDVHEDSLTVVFENNWQPERQVPFNEVRLPPPPDIKKEIS 60
```

Supplementary Figure 3

(c)

```
                     60-62
FXR2Pzebrafish    50 PPPTDYHKDICEGDEVEVYSRANEQEPCGWWLARVRMMKGEFYVIEYAACDATYNEIVTS 109
FXR2Popossum      60 PPPADYSKEITEGDEVEVYSRANEQEPCGWWLARVRMMKGDFYVIEYAACDATYNEIVTL 119
FXR2Pmouse        60 PPPADYNKEITEGDEVEVYSRANEQEPCGWWLARVRMMKGDFYVIEYAACDATYNEIVTL 119
FXR2Pchimpanzee   60 PPPADYNKEITEGDEVEVYSRANEQEPCGWWLARVRMMKGDFYVIEYAACDATYNEIVTL 119
FXR2Phuman        60 PPPADYNKEITEGDEVEVYSRANEQEPCGWWLARVRMMKGDFYVIEYAACDATYNEIVTL 119
                     ***:** *:* *****************************:******************

                                                                       394-395
FXR2Pzebrafish   350 PFIFVGTKENISNAQALLEYHVAYLQEVEQLRLERLQIDEQLRQIGVGYRAPPSRSGSGV 409
FXR2Popossum     344 PFIFVGTRENISNAQALLEYHLSYLQEVEQLRLERLQIDEQLRQIGLGFRTPGSGRGNSS 403
FXR2Pmouse       344 PFIFVGTRENISNAQALLEYHLSYLQEVEQLRLERLQIDEQLRQIGLGFRPPGSGRGGSG 403
FXR2Pchimpanzee  344 PFIFVGTRENISNAQALLEYHLSYLQEVEQLRLERLQIDEQLRQIGLGFRPPGSGRGSGG 403
FXR2Phuman       344 PFIFVGTRENISNAQALLEYHLSYLQEVEQLRLERLQIDEQLRQIGLGFRPPGSGRGSGG 403
                     *******.*************::***********************:*:*.* *  *..

                               473-474            492-494
FXR2Pzebrafish   465 DRESRPKGVGADDRGSKRGGRGRGSSAGRGRGG-PGPR-----NINTISSVLRDPDSNPY 518
FXR2Popossum     456 REEPDRPGHGDQDTSSRGEEGRRRPMGGRGRGSNPAPRLPSKYTTSSISSVLKDPDSNPY 515
FXR2Pmouse       462 REESNRAGPGDRDPPSRGEESRRRPIGGRGRGPPPVPRPTSRYNSSSISSVLKDPDSNPY 521
FXR2Pchimpanzee  460 REEPNRAGPGDRDPPTRGEESRRRPTGGRGRGPPPAPRPTSRYNSSSISSVLKDPDSNPY 519
FXR2Phuman       460 REEPNRAGPGDRDPPTRGEESRRRPTGGRGRGPPPAPRPTSRYNSSSISSVLKDPDSNPY 519
                       *.   * *  *  :.     * . .*****  * **     . .:*****.*******

                           529-530  538-540
FXR2Pzebrafish   519 SLLE-GEGEQGGDTDASESMGGIDRRRRSRRRRNDLEPSLMDAAANESDGQGATSENGLE 577
FXR2Popossum     516 SLLDTSEPEPPVDSEPGEPPPASARRRRSRRRRTDEDRTIIDGGL-ESDGPNLA-ENGLE 573
FXR2Pmouse       522 SLLDTSEPEPPVDSEPGEPPPASARRRRSRRRRTDEDRTVMDGGL-ESDGPNMT-ENGLE 579
FXR2Pchimpanzee  520 SLLDTSEPEPPVDSEPGEPPPASARRRRSRRRRTDEDRTVMDGGL-ESDGPNMT-ENGLE 577
FXR2Phuman       520 SLLDTSEPEPPVDSEPGEPPPASARRRRSRRRRTDEDRTVMDGGL-ESDGPNMT-ENGLE 577
                     ***: .* *   *::..*.  .  *********.* : :::*..  **** . : *****

                                                                      626-627
FXR2Pzebrafish   578 EEGRPQRRNRSRRRRNRANRPEGGSTSRDRQPVTVADFISRAESQSRQNLPGKEAN-PHV 636
FXR2Popossum     574 EESKPQRRNRSRRRRNRGNRAD-GSISRDRQPVTVADYISRAESQSRQRQPPEPTHVPSE 632
FXR2Pmouse       580 DESRPQRRNRSRRRRNRGNRTD-GSISGDRQPVTVADYISRAESQSRQR-PLERTK-PSE 636
FXR2Pchimpanzee  578 DESRPQRRNRSRRRRNRGNRTD-GSISGDRQPVTVADYISRAESQSRQRPPLERTK-PSE 635
FXR2Phuman       578 DESRPQRRNRSRRRRNRGNRTD-GSISGDRQPVTVADYISRAESQSRQRPPLERTK-PSE 635
                     :*..*************.**.: ** * *********:**********. * : :: *
```

Supplementary Figure 4

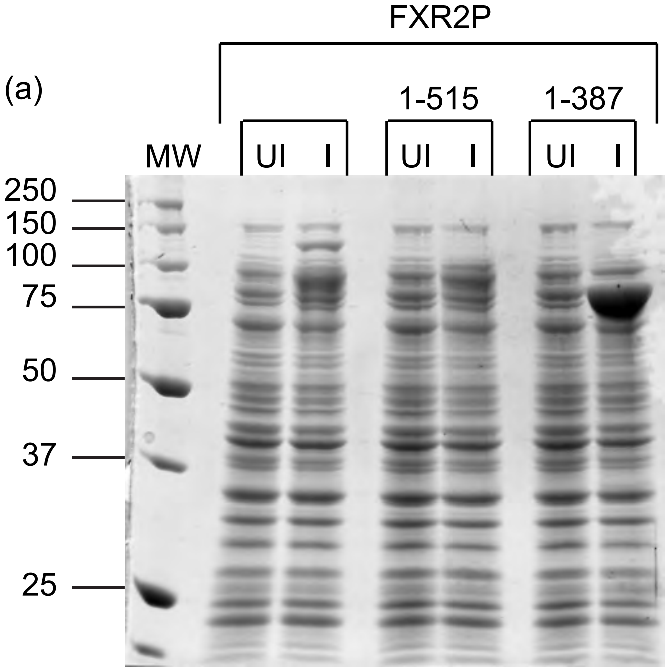


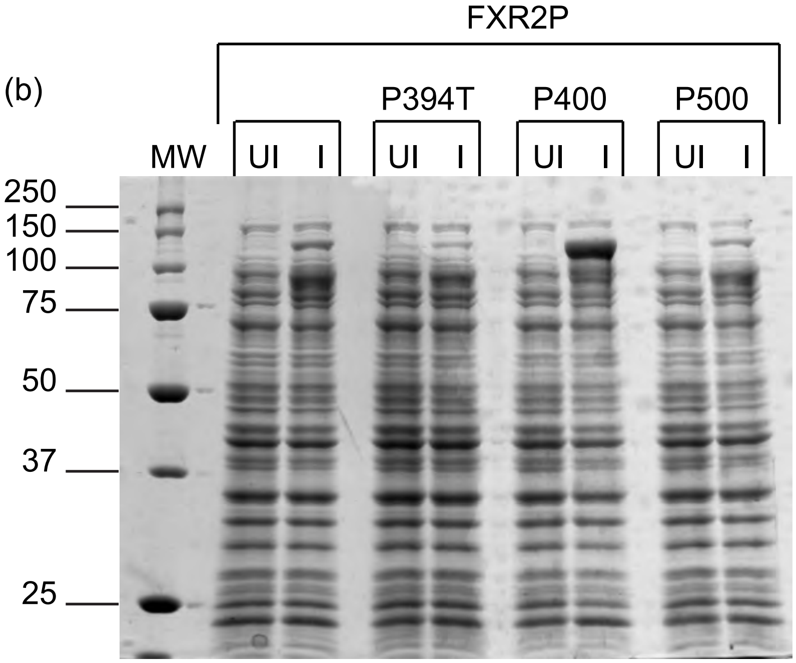


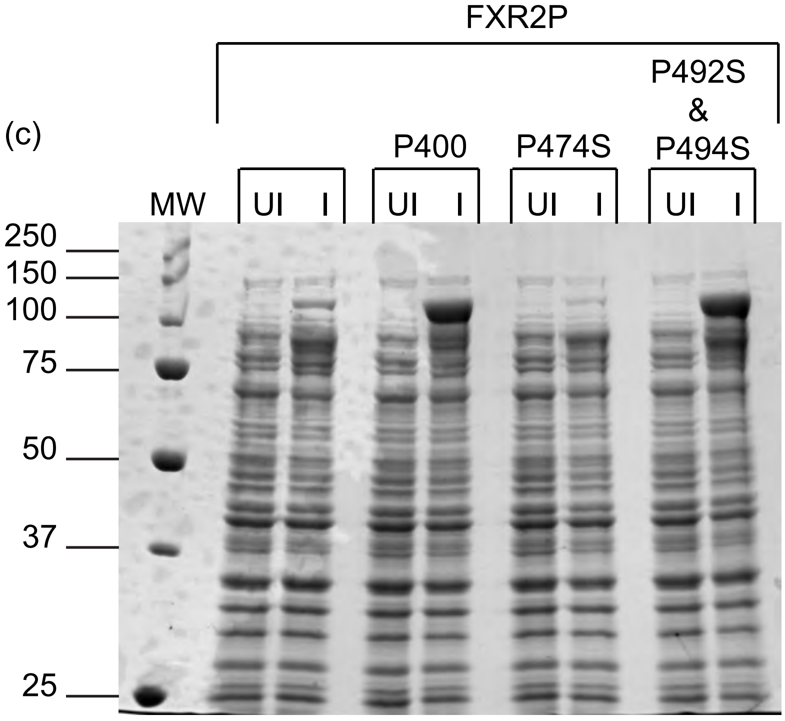

Supplementary Figure 5

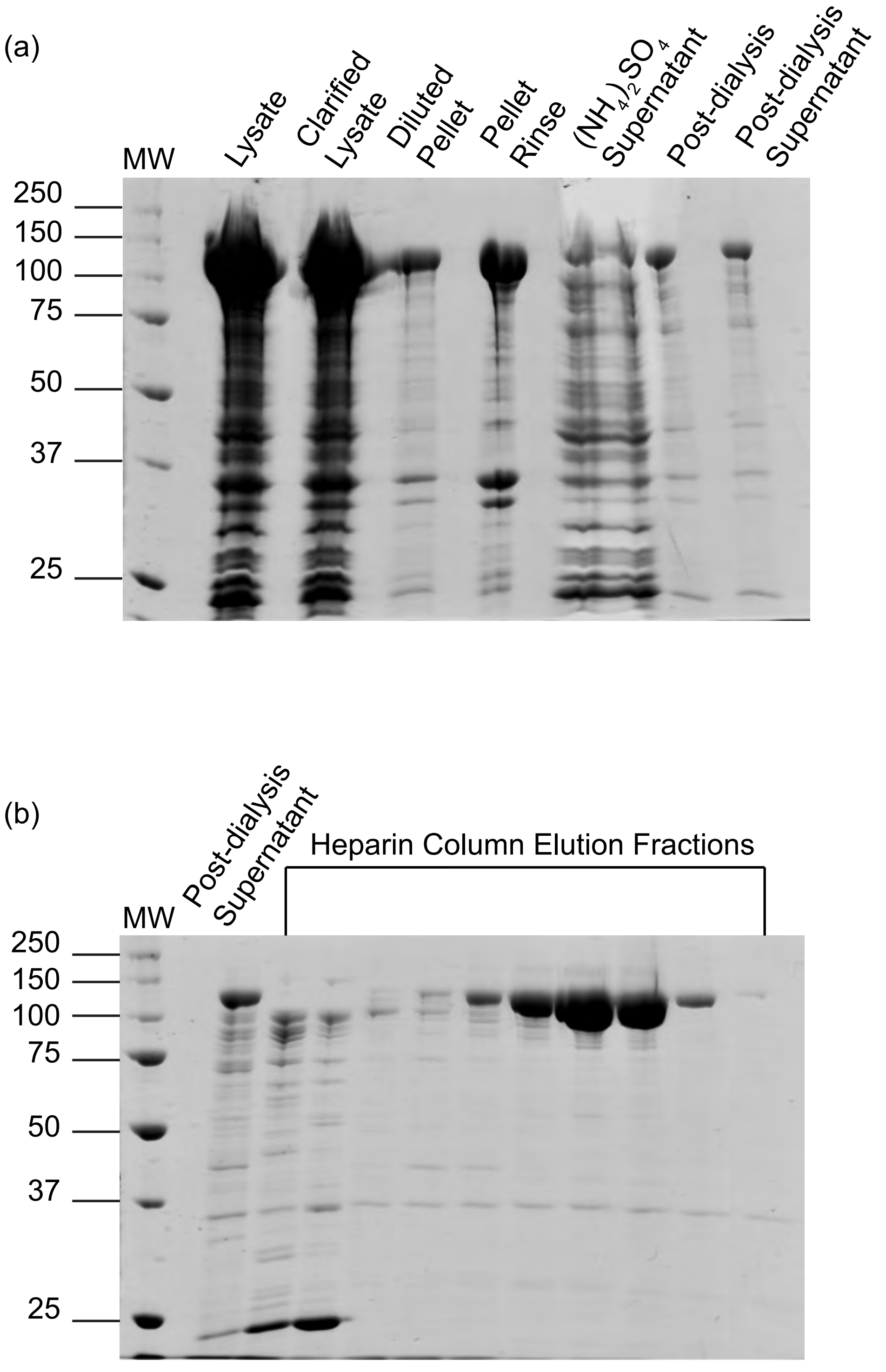

Supplementary Figure 6

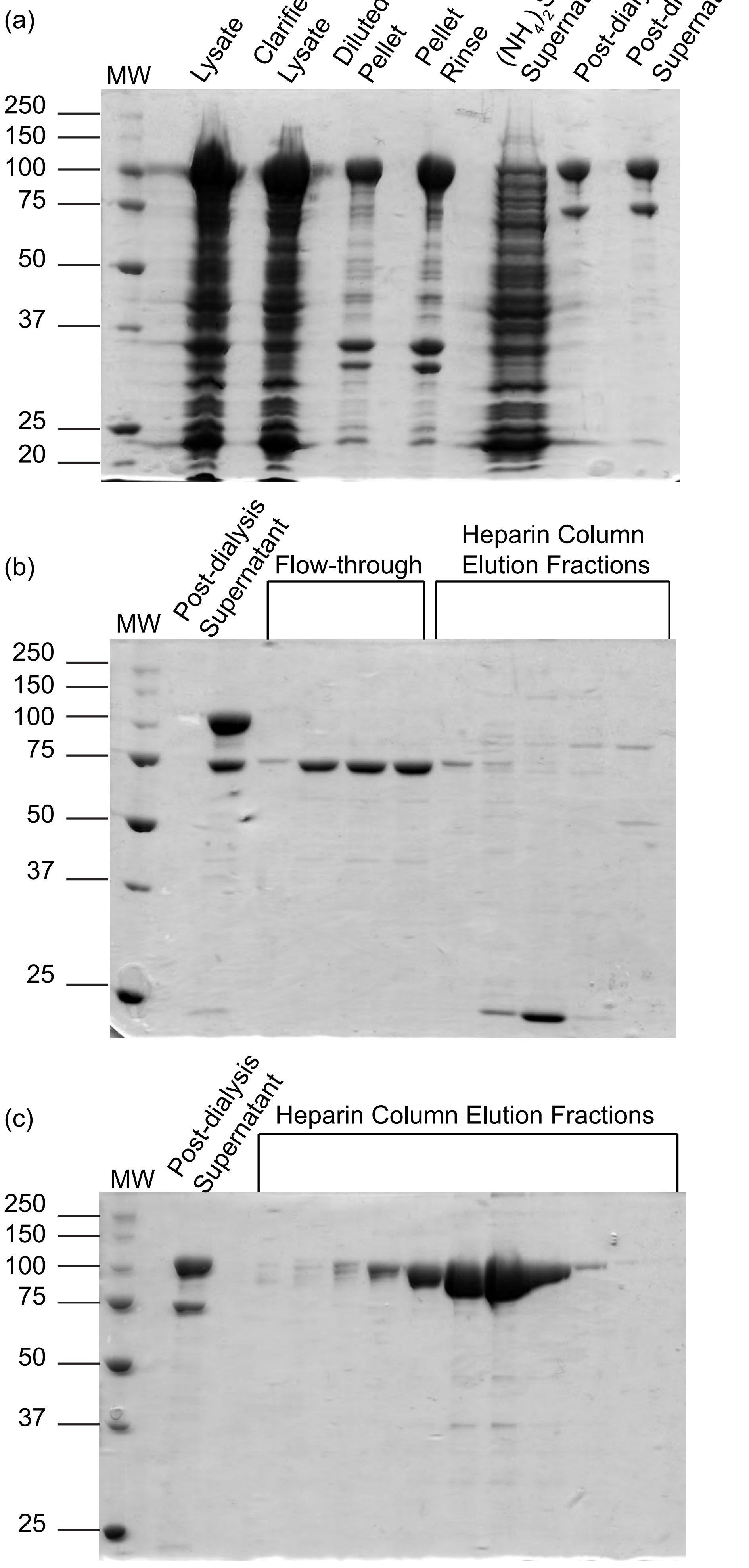

Supplementary Figure 7

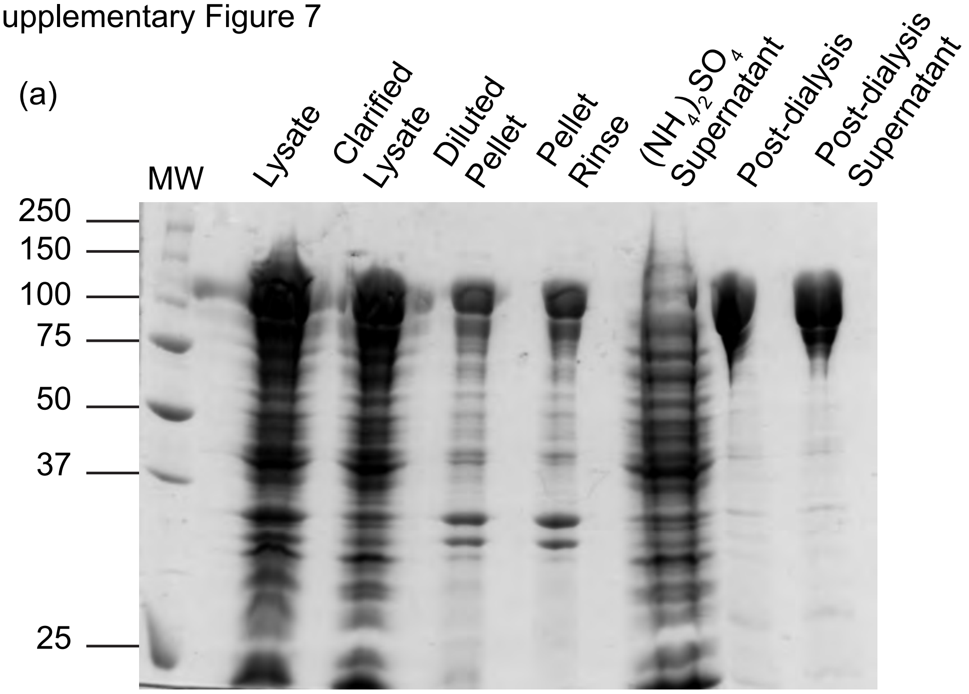


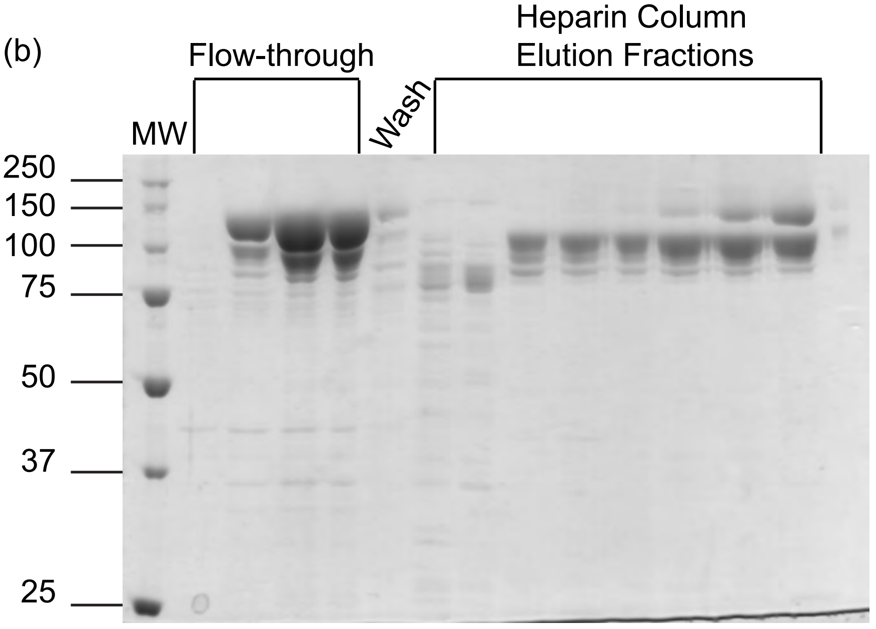

Supplementary Figure 7

(c)

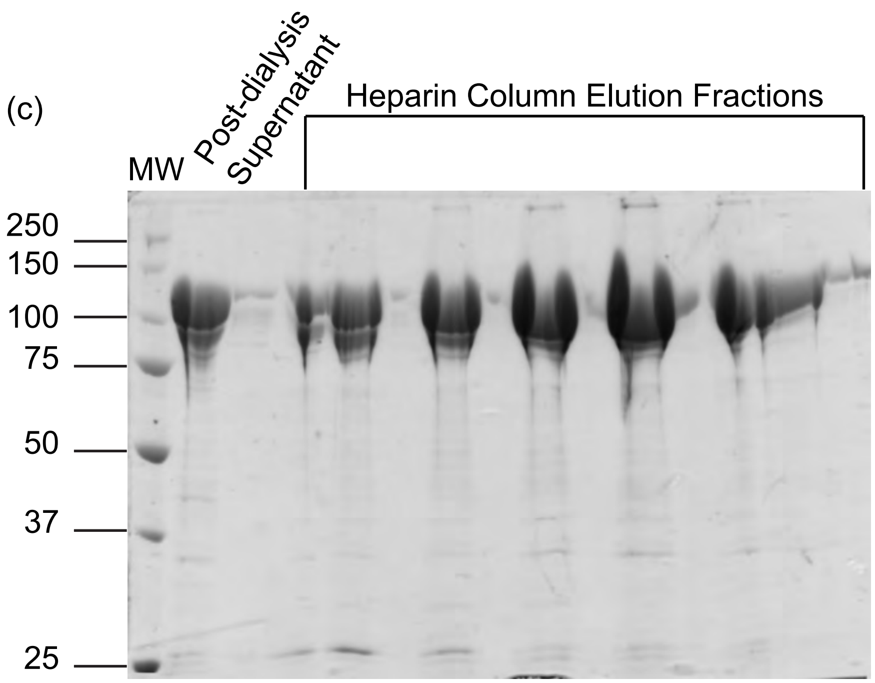


(d)

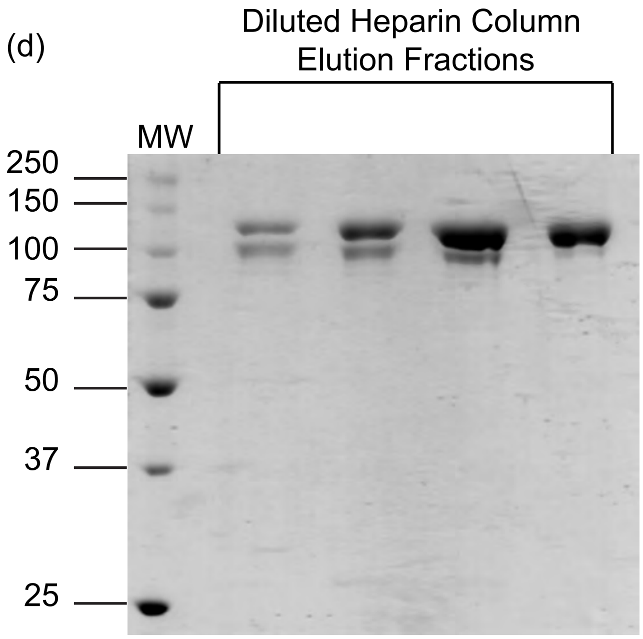

Supplementary Figure 8

(a)

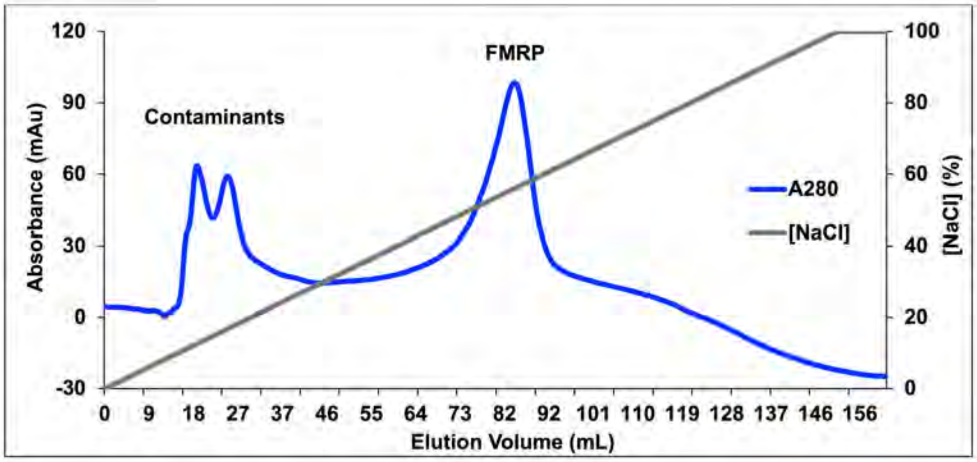


(b)

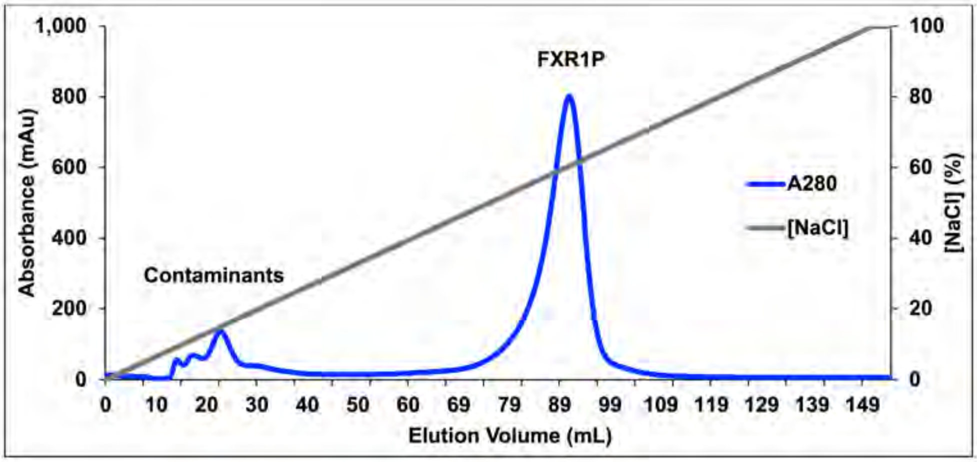


(c)

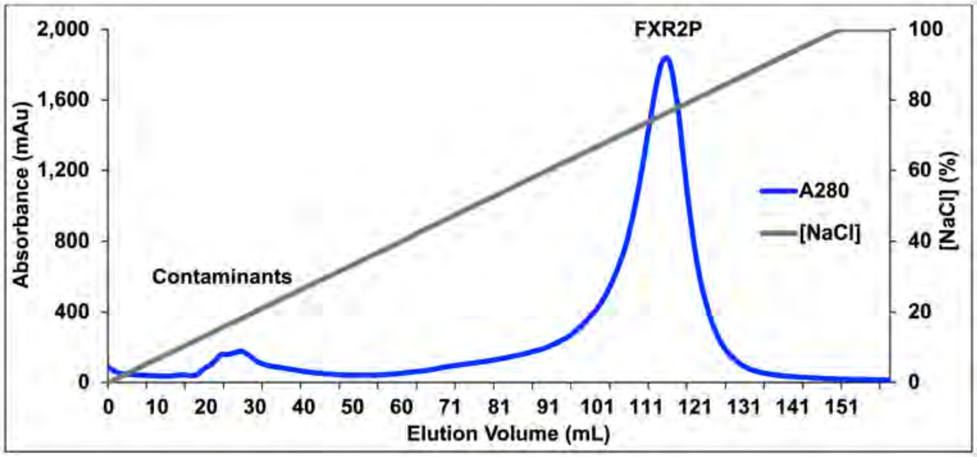
